# A V0-ATPase-dependent apical trafficking pathway maintains the polarity of the intestinal absorptive membrane

**DOI:** 10.1101/412122

**Authors:** Aurélien Bidaud-Meynard, Ophélie Nicolle, Markus Heck, Grégoire Michaux

## Abstract

Intestine function relies on the strong polarity of intestinal epithelial cells and the array of microvilli forming a brush border at their luminal pole. Combining genetic RNAi screen and *in vivo* super-resolution imaging in the *C*. *elegans* intestine, we uncovered that the V0 sector of the V-ATPase (V0-ATPase) controls a late apical trafficking step, involving RAB-11 endosomes and the SNARE SNAP-29, necessary to maintain the polarized localization of both apical polarity modules and brush border proteins. We show that the V0-ATPase pathway also genetically interacts with glycosphingolipids in enterocyte polarity maintenance. Finally, we demonstrate that depletion of the V0-ATPase fully recapitulates the severe structural, polarity and trafficking defects observed in enterocytes from patients with Microvillus inclusion disease (MVID) and used this new *in vivo* MVID model to follow the dynamics of microvillus inclusions. Hence, we describe a new function for the V0-ATPase in apical trafficking and epithelial polarity maintenance and the promising use of *C*. *elegans* intestine as an *in vivo* model to better understand the molecular mechanisms of rare genetic enteropathies.

**Summary statement:** V0-ATPase controls a late apical trafficking step involved in the maintenance of the apical absorptive intestinal membrane and its depletion phenocopies the trafficking and structural defects of MVID in *C*. *elegans*.

## Introduction

Considering that almost all nutrients in the diet are absorbed into blood across the epithelial layer forming the small and large intestinal mucosa (Kiela and Ghishan, 2016), proper establishment and maintenance of both the strong polarity of enterocytes and the array of microvilli forming an absorptive brush border at their apical plasma membrane (PM) are essential to ensure intestinal functions. In most species, including *C*. *elegans*, epithelial cell polarity is achieved by actin and microtubule cytoskeleton reorganization, trafficking and signalling-mediated polarized targeting of the CDC-42/PAR-3/PAR-6/PKC-3 (CDC-42/PAR) and Crumbs/PALS1/Patj (CRB) modules at the apical pole and the Dlg/Scribble/Lgl (SCRIB) module at the basolateral pole as well as domain-specific lipid distribution (Rodriguez-Boulan and Macara, 2014). Particularly, CDC-42 has been shown to be a major player of polarity maintenance by controlling the apical localization of PAR module components in *C*. *elegans* intestine (Shafaq-Zadah et al., 2012) and MDCK cysts (Martin-Belmonte et al., 2007). The establishment of the brush border, an extended PM surface for efficient nutrient uptake, is thought to be highly dependent on actin cytoskeleton regulating factors which create the force necessary for the onset and maintenance of the microvilli (Heintzelman and Mooseker, 1990a, Heintzelman and Mooseker, 1990b, Crawley et al., 2014, Saegusa et al., 2014).

Polarity establishment and brush border formation are early events in intestine development that both rely on related cytoskeletal and trafficking pathways. In *C*. *elegans*, Clathrin and AP-1 (Shafaq-Zadah et al., 2012, Zhang et al., 2012), glycosphingolipids (GSLs) (Zhang et al., 2011), and the kinase Lats/WTS-1 (Kang et al., 2009) have been shown to play a major role in the sorting of polarity modules and luminogenesis. In mammalian intestine cellular models, the trafficking proteins Annexin 2, STX3, Rab8a and Rab11a, the actin binding proteins Drebin E, Myosin 6 and Myosin 5 have been involved in the polarized localization of PM transporters and brush border integrity (Hegan et al., 2012, Vogel et al., 2015, Vacca et al., 2014, Hein et al., 2011) while the latter has been also involved in the apical confinement of the PAR polarity module (Michaux et al., 2016).

Particularly, these studies highlighted the central role of RAB-11-dependent apical trafficking in polarity maintenance and lumen integrity. Indeed, studies in MDCK cells demonstrated the critical function of RAB-11^+^ endosomes in apical PM and lumen generation by mediating PAR-3 targeting and Cdc42 activation through the exocyst complex (Colombie et al., 2017, Bryant et al., 2010) as well as the adverse effect of dominant-negative Rab11 (Rab11^DN^) expression on the apical localization of phospho-ezrin and E-Cadherin (Desclozeaux et al., 2008). Furthermore, expression of Rab-11^DN^ or depletion of the Rab11^+^ endosomes-associated motor Myosin 5B (Myo5B) decreased both ezrin phosphorylation at the apical PM and microvilli size in intestinal cells model (Dhekne et al., 2014), and these polarity and structural consequences were confirmed in Rab11a intestinal-specific knock out mice model (Sobajima et al., 2014). Finally, defective RAB-11-dependent trafficking has been associated with congenital enteropathies, which are characterized by polarity and brush border structural defects (Vogel et al., 2017a). However, little is known about the relationship between polarity modules, brush border and apical trafficking components and their sequential involvement in establishing and maintaining an absorptive intestinal apical PM, especially *in vivo*.

*C*. *elegans* intestine is composed of 20 polarized enterocytes similar to that of mammalian cells and the nematode embryonic and larval stages have been widely used to understand polarity and lumen establishment and maintenance, respectively (McGhee, 2007). By performing an RNAi screen to target a library of trafficking and cytoskeleton factors in *C*. *elegans* larvae, we unveiled that the depletion of the V0 sector of the vacuolar-ATPase (V-ATPase) induces a cytoplasmic and basolateral mislocalization of both PAR polarity module and brush border components. The V-ATPase is a large hetero-multimeric complex (20 subunits in *C*. *elegans, vha-1* to *vha-18, spe-5* and *unc-32*), organized in two main sectors: a V1 sector responsible for ATP hydrolysis which provides the energy to the V0 sector, which transports protons (Lee et al., 2010). The whole V-ATPase complex is responsible for the acidification of intracellular compartments, such as endosomes or lysosomes and plays a major role in PM proteins trafficking (Maxson and Grinstein, 2014), while its V0 transmembrane sector (V0-ATPase) has been implicated in the fusion to the yeast vacuole (Baars et al., 2007) as well as that of synaptic vesicles in drosophila and humans (Hiesinger et al., 2005, Di Giovanni et al., 2010), in an acidification-independent manner. In *C*. *elegans*, the V0-ATPase subunits *vha-5, vha-6* and *vha-19* are required for exosome containing vesicle apical exocytosis in the epidermis (Liegeois et al., 2006), intestinal lumen pH regulation (Allman et al., 2009) and RME-2 receptor trafficking in oocytes (Knight et al., 2012) but no role in polarity maintenance has been assigned to this complex before.

Here, we show that interfering with the V0-ATPase-mediated polarity maintenance mechanism in *C*. *elegans* fully recapitulates the polarity, trafficking and structural phenotypes observed in patients suffering from Microvillus inclusion disease (MVID, OMIM: 251850). MVID is caused by mutations in the genes coding for myosin-5B (MYO5B) (Muller et al., 2008), syntaxin-3 (STX3) (Wiegerinck et al., 2014) or Munc18-2/STXBP-2 (STXBP2) (Vogel et al., 2017b), three proteins regulating apical trafficking. The major hallmarks of this disease are a microvillus atrophy, the rare formation of microvillus inclusions, as well as a loss of subapical RAB-11^+^ endosomes (Schneeberger et al., 2018). Hence, we propose that the V0-ATPase is essential to control apical trafficking and polarity maintenance.

## Results

### V0-ATPase depletion affects the polarity of brush border and polarity components in the *C*. *elegans* intestine

To identify new common regulators of polarity and brush border maintenance *in vivo*, we performed a two-step RNAi screen targeting 408 genes essential for survival, conserved in humans and implicated in membrane trafficking and cytoskeleton regulation. First, we examined the effect of their silencing on the localization of the polarity module component CDC-42 at the apical PM of enterocytes of *C*. *elegans* larvae. We then performed a secondary screen to evaluate the ability of the hits decreasing CDC-42 apical localization to also disturb the polarity of the brush border specific actin cross-linker ERM-1 (Table S1 and Fig S1A). This systematic RNAi screen first confirmed the involvement of clathrin (*chc-1*), AP-1 (*aps-1*) and GSLs biosynthesis enzymes (i.e. *sptl-1, fasn-1, pod-2*) (Zhang et al., 2012, Shafaq-Zadah et al., 2012, Zhang et al., 2011) in the maintenance of intestinal polarity (Table S1) and identified several genes belonging to the biosynthetic and endocytic pathways, as well as 10 subunits of the V-ATPase complex, a major player of intracellular trafficking by regulating endosomal acidification (Table S1). This screen performed using overexpressed markers, was then confirmed by silencing several subunits of the V-ATPase complex and studying the endogenous localization of genome-edited mNeongreen or GFP-tagged ERM-1 as well as the polarity module components PAR-6 and PKC-3. Consistently, knockdown of all the V-ATPase complex subunits decreased by 15-40% the apical/cytoplasmic ratio of the PAR module proteins and of ERM-1 (Fig 1A-F). This indicates that a V-ATPase acidification-dependent mechanism is involved in the apical confinement of brush border and polarity module components. Intriguingly, depletion of the subunits *vha-1, −4* and *-19*, belonging to the V0-ATPase, also induced a basolateral localization of ERM-1, while no basolateral mislocalization was observed upon silencing of the V1-ATPase subunits *vha-8*, which has been shown to inhibit V-ATPase-dependent acidification (Ji et al., 2006) and *vha-15* (Fig 1A, G). Furthermore, endogenous PAR-6 and PKC-3 could also be specifically observed at the basolateral PM in ∼20% of the worms upon V0-ATPase subunits depletion (Fig 1B). To evaluate if V0-ATPase silencing induces a global polarity inversion of the enterocytes we studied the effect of the silencing of the V0-ATPase subunit *vha-1*, which destabilizes the whole V0-ATPase sector (Fig S1 B-C) on the polarized localization of the apical peptide transporter PEPT-1 (OPT-2) and the P-glycoprotein PGP-1 as well as the basolateral pyruvate transporter SLCF-1. Notably, silencing of *vha-1* did not induce a significant effect on the cortical confinement of those highly polarized proteins and none of them accumulated at the contralateral PM (Fig 1H-K and Fig S2A), supporting the notion that the enterocyte polarity is only partially compromised upon V0-ATPase depletion. Overall, these data indicate that the V0-ATPase sector specifically controls the polarized localization of both brush border (ERM-1) and polarity module components (PAR-6, PKC-3) in enterocytes.

**Figure 1.**
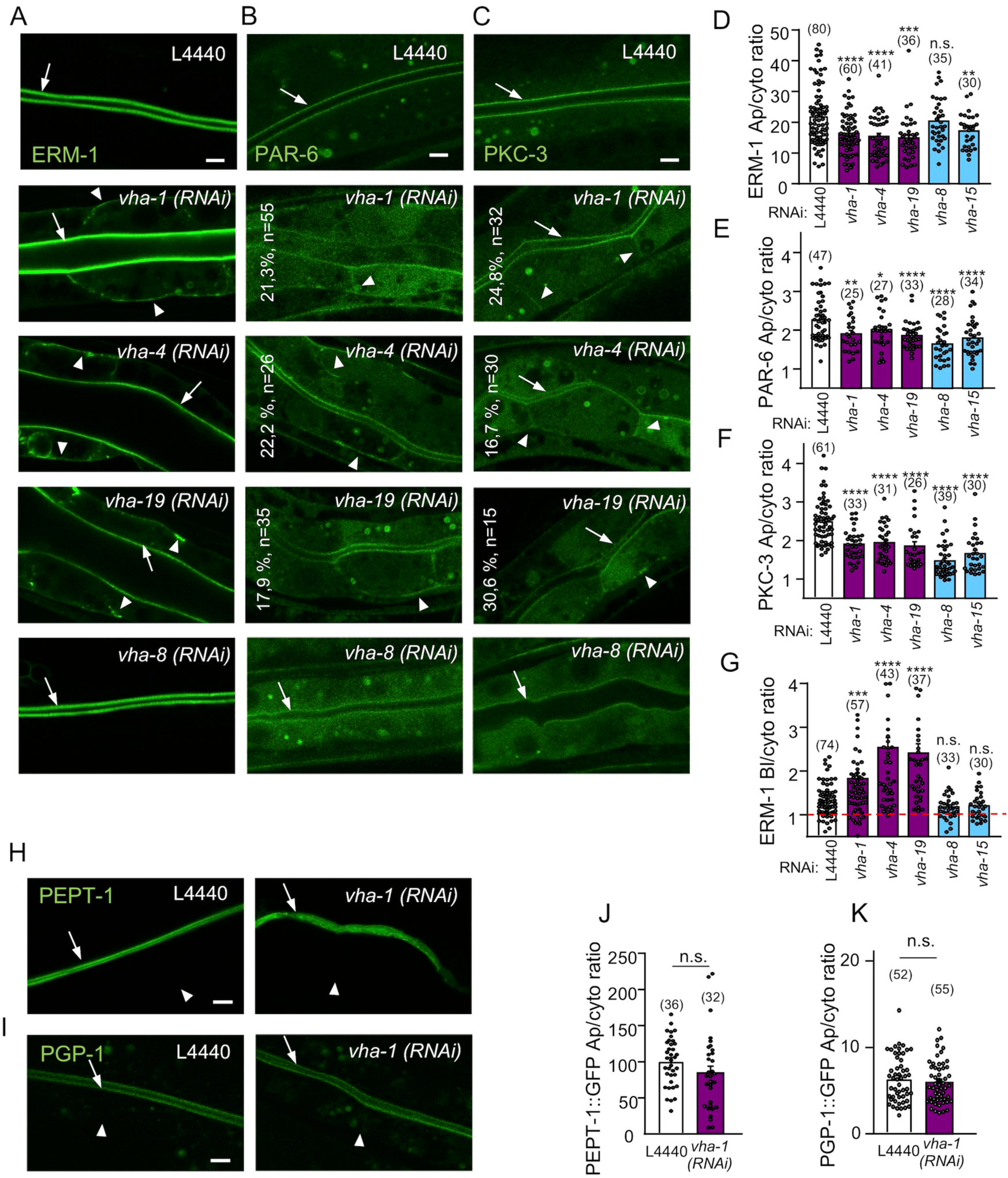
V0-ATPase controls the apical localization of polarity and brush border components. (A-C) Larvae expressing endogenous ERM-1::mNG, PAR-6::GFP and GFP::PKC-3 were imaged after 72h silencing of V0-ATPase (*vha-1(RNAi))* or V1-ATPase (*vha-8(RNAi))* subunits. Arrows show the intestinal apical PM. Arrowheads show the basolateral mislocalization of markers upon *vha-1(RNAi)*. The percentage of worms displaying basolateral PAR polarity markers and the number of worms analyzed are indicated on each panel. (D-F) Quantification of the apical/cytoplasmic (Ap/cyto) ratio of ERM-1::mNG (D), PAR-6::GFP (E) and GFP::PKC-3 (F) upon 72h silencing of V0-ATPase (*vha-1, −4, −19*) or V1-ATPase (*vha-8, −15*) subunits (n=3-5 independent experiments). (G) V0-ATPase subunits (*vha-1, −4, −19*) knockdown specifically induces a basolateral relocalization of ERM-1. Histogram shows the basolateral/cytoplasmic (Bl/cyto) ratio of ERM-1::mNG. Dotted red line indicates a ratio of 1, which means no basolateral localization. (H-K) V0-ATPase silencing does not affect the polarity of the apical transmembrane proteins PEPT-1 and PGP-1. Strains expressing exogenously (PEPT-1) or endogenously (PGP-1)-tagged markers were silenced for *vha-1* or *vha-8* during 72h and imaged. Histograms in (J-K) represent the quantification of the Ap/cyto ratio. Arrows show the intestinal apical PM. Arrowheads show the basolateral mislocalization of markers upon *vha-1(RNAi)*. Histograms are mean ± SEM, dots represent individual worms and total number of worms from 3 independent experiments is indicated in brackets. n.s., non-significant, *p<0.05, **p<0.01, ***p<0.001, ****p<0.0001. Scale bars, 5 μm.

### V0-ATPase depletion specifically affects the apical recycling pathway

The V-ATPase complex controls the acidification of various endosomes along the biosynthetic and endocytic-recycling routes (Forgac, 2007). Using Transmission Electron Microscopy (TEM), we observed the presence of abnormal large organelles (0.1–0.3 μm in diameter with variable electron density and heterogeneous content) with features of vesicles and lysosomes upon inhibition of both V0 and V1-ATPase subunits (Fig 2A-B). These “mixed” organelles have been already described in the *C*. *elegans* epidermis and in mammalian cells upon V0-/V1-ATPase silencing or chemical inhibition of the V-ATPase complex function, and have been proposed to affect protein sorting along the secretory and lysosomal pathways (Sobota et al., 2009, Liegeois et al., 2006), probably due to an endosome acidification failure. The presence of mixed organelles was confirmed by the accumulation of several endosomal markers in large cytoplasmic structures, such as LMP-1 (lysosomes), RAB-7 (late endosomes) and RAB-10 (basolateral recycling endosomes) (Fig 2C and S2B), upon both V0-and V1-ATPase silencing.

**Figure 2.**
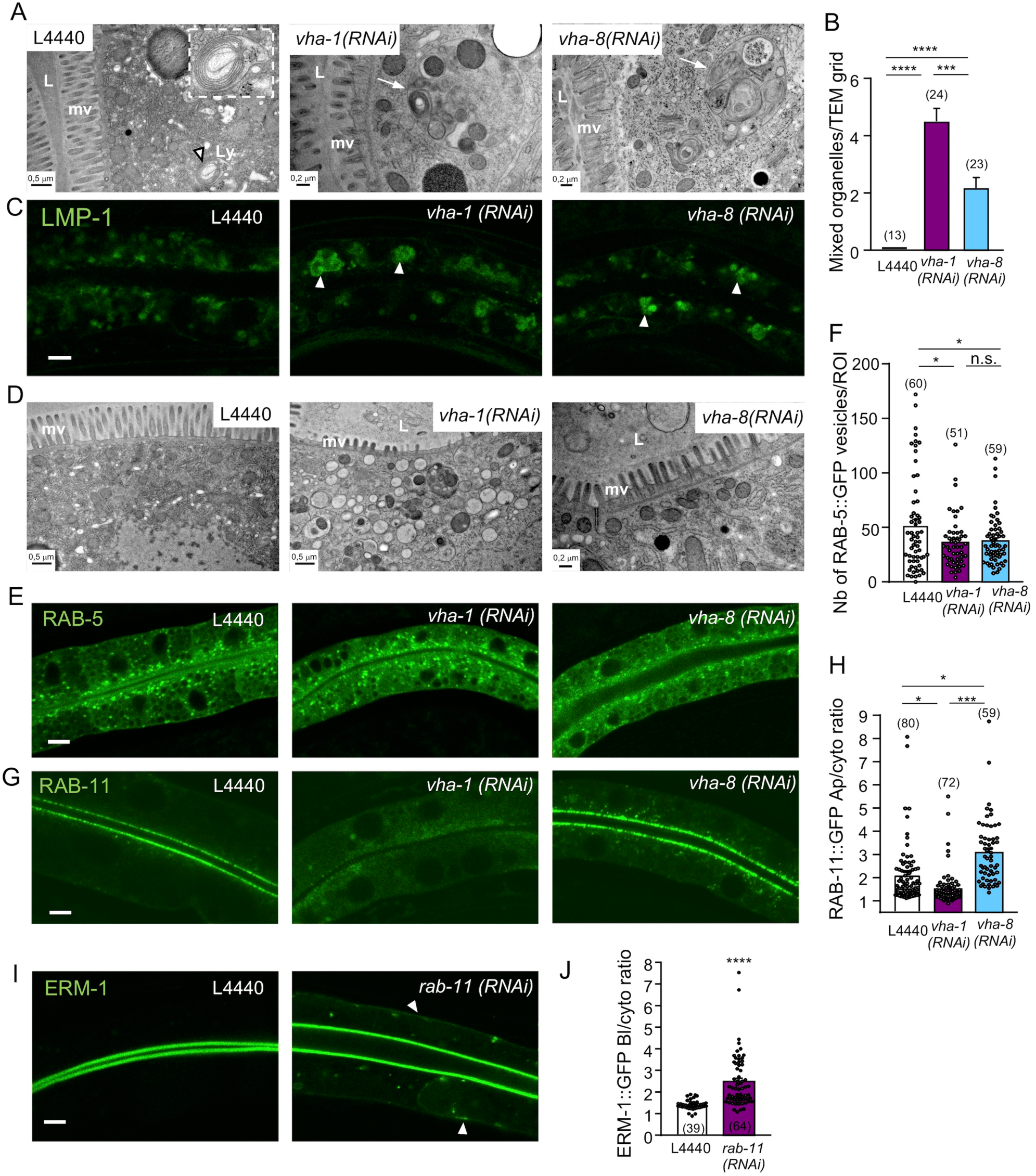
V0-ATPase affects apical trafficking via RAB-11 apical recycling endosomes. (A-C) V-ATPase complex silencing affects the morphology of lysosomes. (A) Ultrastructural characterization of transversely-sectioned control (L4440), V0-ATPase (*vha-1*) or V1-ATPase (*vha-8*) depleted worms (72h RNAi). While lysosomes with normal morphology and size (∼0.5μm) were rarely observed in control worms (left panel, arrowhead and insert), cytoplasmic enlarged “mixed organelles” with distinct lysosomal and vesicular regions were observed upon both *vha-1* and *vha-8(RNAi)* (arrows). (B) Quantification of the number of mixed organelles (n=1-5grid/larvae, n=7-13 larvae). (C) shows the accumulation of the lysosomal marker LMP-1::GFP in large cytoplasmic structures (arrowheads) upon both *vha-1* and *vha-8(RNAi)*. (D) Ultrastructural characterization of transversely-sectioned control (L4440), V0-ATPase (*vha-1*) and V1-ATPase (*vha-8*) depleted worms. Middle panel shows a specific and massive accumulation of cytoplasmic electron-lucent vesicles upon *vha-1(RNAi)*. (E-H) V0- or V1-ATPase silencing marginally decreases the number of RAB-5^+^ vesicles but V0-ATPase knockdown specifically disrupts RAB-11^+^ apical recycling endosomes. Confocal imaging of the endosomal markers of apical early/sorting (RAB-5, E) and apical recycling (RAB-11, G) endosomes in control, *vha-1(RNAi)* and *vha-8(RNAi)* conditions. (F-H) Quantification of the number of RAB-5+ vesicles (F) and RAB-11 apical/cytoplasmic (Ap/cyto) ratio (H) (n=3). (I-J) *rab-11(RNAi)* phenocopies V0-ATPase depletion on ERM-1::GFP polarity. ERM-1::GFP expressing worms were silenced for *rab-11* during 72h and imaged. Arrowheads show the basolateral accumulation of ERM-1. (J) shows ERM-1 basolateral/cytoplasmic (Bl/cyto) ratio quantification (n=3). Histograms are mean ± SEM on each panel, dots represent individual worms and the total number of worms from 3 independent experiments is indicated in brackets. L, lumen; mv, microvilli; Ly, lysosome. n.s., non-significant, *p<0.05, ***p<0.001, ****p<0.0001. Scale bars, 0.2 or 0.5µm (A, D) and 5 μm (C, E, G, I).

Strikingly, V0-but not V1-ATPase depletion led to the specific accumulation of electron-lucent vesicular structures in the cytoplasm (Fig 2D), which were reminiscent of an apical trafficking defect. The number of RAB-5^+^ endosomes was slightly decreased upon both V0 and V1-ATPase silencing (Fig 2E-F). However, RAB-11 endosomes virtually disappeared from the apical PM and accumulated in the cytoplasm upon V0-but not V1-ATPase knockdown (Fig 2G-H). Considering that *rab-11* knockdown also induced a basolateral mislocalization of ERM-1 (Fig 2I-J), similarly to *vha-1(RNAi)* (Fig 1B, G) this indicates that V0-ATPase controls the enterocyte polarity by regulating a late trafficking step through RAB-11^+^ endosomes.

### SNAP-29 is involved in V0-ATPase-mediated polarity maintenance

To identify other molecular components of this V0-ATPase-mediated control of intestinal polarity beyond the interaction with RAB-11, the effect of V0- or V1-ATPase depletion was assayed on several soluble trafficking factors with a highly polarized localization. The PM localization of CHC-1/clathrin and the syntaxin UNC-64 (the presumptive mammalian STX3 ortholog) was strongly decreased and increased, respectively (Fig 3 A-B) while that of the syntaxin SYX-1 (SYN-1) was only slightly decreased (Fig S2C) upon all V-ATPase complex subunits depletion, suggesting a trafficking defect, but no sector-specificity. Strikingly, *vha-1/V0-ATPase(RNAi)* specifically decreased the apical localization of the soluble N-ethylmaleimide-sensitive factor-attachment protein receptors (SNAREs) component SNAP-29, while *vha-8/V1-ATPase(RNAi)* did not (Fig 3C). SNAP-29 belongs to the SNAP23/25/29 family of SNARE proteins that interact with syntaxins and sec/munc proteins to promote vesicle fusion (Sudhof and Rothman, 2009). Its depletion has been shown to affect the morphology of various organelles, including recycling endosomes, and consequently endocytosis, secretion as well as autophagy pathways in *C*. *elegans* intestine (Sato et al., 2011, Guo et al., 2014). To test whether SNAP-29 is involved in polarity maintenance, *snap-29* was silenced by RNAi (Fig S2 D-E) and the PM localization of ERM-1 was assessed. *snap-29* silencing induced a basolateral mislocalization of ERM-1 (Bl/cyto ratio >1.5) in 44.3% of the worms whereas only 20.8% of control (L4440) worms displayed a ratio >1.5 (Fig 3D-E). Finally, the effect of *snap-29(RNAi)* was assayed on a strain co-expressing RAB-11::GFP and the V0-ATPase subunit VHA-6::mCh. While *snap-29* knockdown also induced a dramatic loss of RAB-11 signal, it did not affect that of V0-ATPase (Fig 3 F-G). Hence, these data strongly suggest that SNAP-29 acts downstream of the V0-ATPase to maintain the enterocyte polarity.

**Figure 3.**
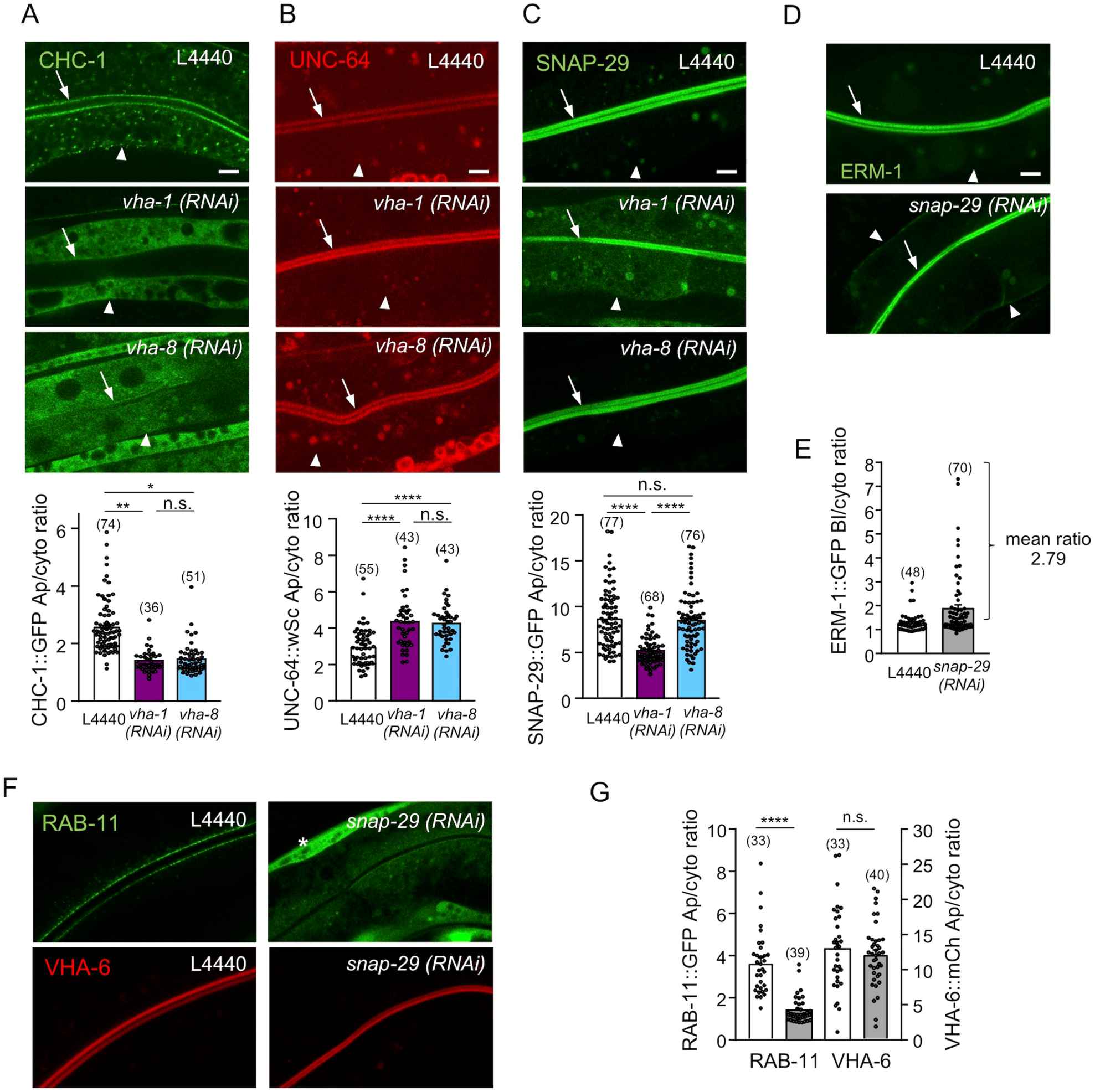
SNAP-29 acts downstream of V0-ATPase in polarity maintenance. (A-C) V0-ATPase silencing specifically decreases the apical localization of SNAP-29::GFP. Strains expressing the indicated markers were silenced for V0-(*vha-1*) or V1-(*vha-8*) ATPase during 72h and imaged. Histograms represent the quantification of the apical/cytoplasmic (Ap/cyto) ratio of each marker. (D-E) *snap-29(RNAi)* induces a basolateral localization of ERM-1. *snap-29* was knocked down by RNAi in ERM-1::GFP-expressing strains and its localization was recorded after 72h. (E) represents the quantification of the basolateral/cytoplasmic (Bl/cyto) ratio of ERM-1::GFP. (F-G) *snap-29(RNAi)* also induces a RAB-11 endosomes loss but does not affect VHA-6 apical localization. *snap-29* was depleted by RNAi in a strain co-expressing RAB-11::GFP and VHA-6::mCh. Arrows and arrowheads indicate the apical and basolateral PM, respectively. The asterisk shows muscular Myo-3P::GFP co-expressed with VHA-6::mCh. (G) shows the quantification of RAB-11::GFP (left panel) and VHA-6::mCh (right panel) apical/cytoplasmic ratio (Ap/cyto). Histograms are mean ± SEM on each panel, dots represent individual worms and the total number of worms from 3 (A-C, E) and 2 (G) independent experiments is indicated in brackets. n.s., non-significant, *p<0.05, **p<0.01, ****p<0.0001. Scale bars, 5 μm.

### High-resolution map of the apical PM revealed that V0-ATPase localizes to microvilli

The V0-ATPase has been localized at the apical PM of intestinal epithelia in *C*. *elegans* (Zhu et al., 2015) and proposed to traffic to the brush border in Caco-2 cells (Collaco et al., 2013). However, conventional confocal imaging in *C*. *elegans* only allowed to determine the apical polarity of markers, but not their precise position regarding the apical PM (e.g. microvilli, terminal web, subapical). The precise localization of the V0-ATPase as well as that of some of the above-mentioned polarity markers was thus assessed by live super-resolution confocal microscopy. This technique confirmed that ERM-1 accumulates in the microvilli, while ACT-5, PGP-1 and SNAP-29 were present at both the microvilli and the underlying terminal web (Fig 4A-B). SNAP-29 has been previously shown to localize on RAB-11^+^ endosomes in *C*. *elegans* intestine (Kang et al., 2011), and this interaction seems to take place at the presumptive terminal web level, where the PAR component PKC-3 mostly accumulates (Fig 4A-D). Finally, the V0-ATPase subunit VHA-6 was localized to the microvilli, above the RAB-11 apical staining (Fig 4D). Overall, these results strongly suggest that the V0-ATPase localizes at microvilli from where it controls the apical localization of brush border and polarity modules by regulating a late trafficking step through RAB-11^+^ endosomes in a SNAP-29-dependent manner.

**Figure 4.**
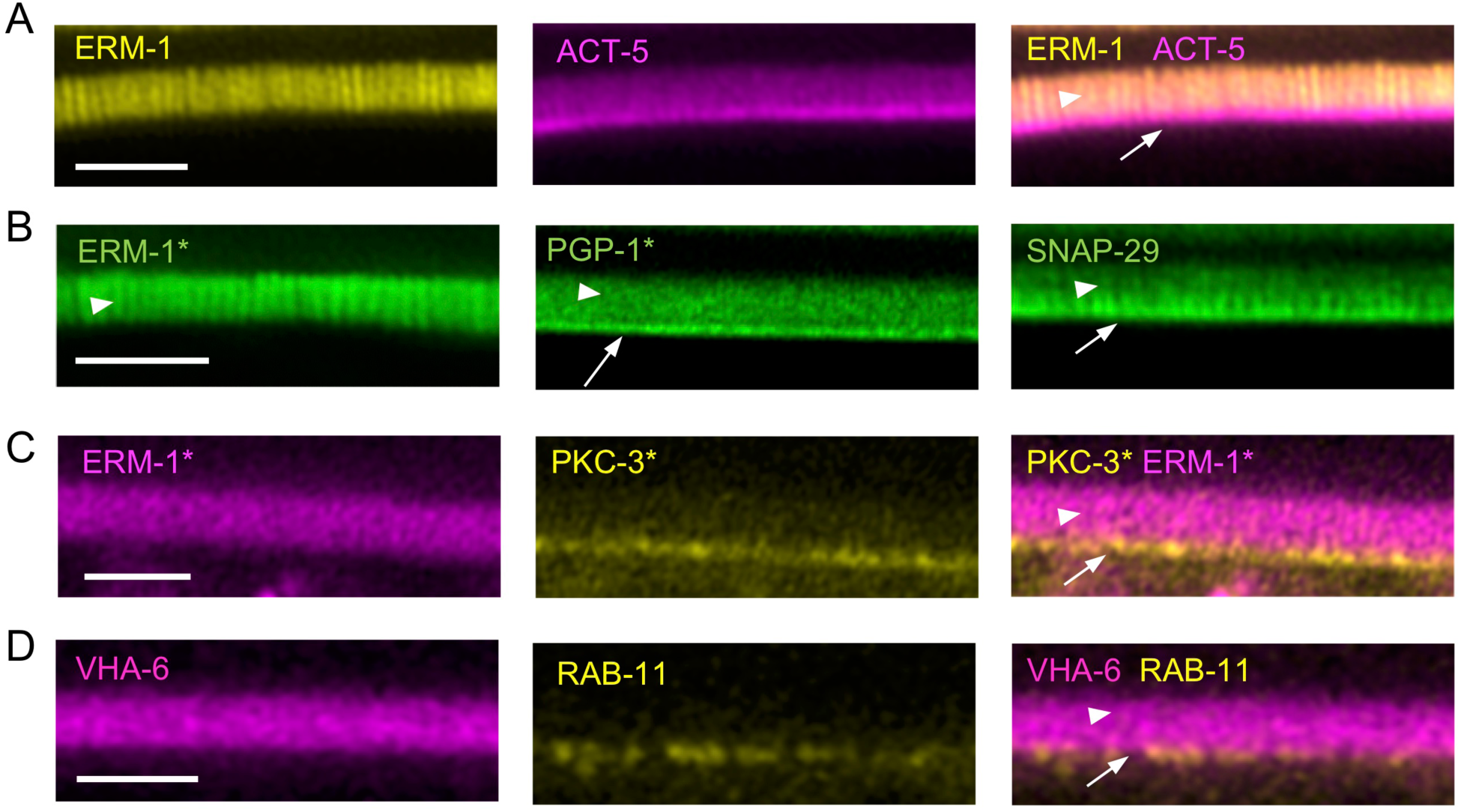
Super-resolution imaging of the intestinal apical PM. (A-D) Strains expressing the indicated markers were mounted and imaged. * indicates endogenously-tagged markers, arrowheads and arrows indicate microvilli and presumptive terminal web, respectively. Scale bars, 2 μm.

### V0-ATPase interacts with GSLs but not with AP-1 in intestinal polarity maintenance

Interestingly, two trafficking and possibly cooperating pathways have been invoked as major regulators of intestinal polarity maintenance in *C*. *elegans*: GSLs and Clathrin/Clathrin adaptor AP-1. Particularly, depletion of the AP-1 complex subunit *aps-1* or of enzymes along the GSLs biosynthetic pathway (i.e. *let-767, acs-1, pod-2* or *sptl-1*) has been shown to induce the contralateral PM localization of polarized proteins and a loss of RAB-11 endosomes, similarly to *vha-1* depletion (Shafaq-Zadah et al., 2012, Zhang et al., 2012, Zhang et al., 2011). First, to test the putative genetic interaction between the V0-ATPase and the GSLs and AP-1 pathways, *vha-1* and *aps-1* were co-depleted by RNAi and *vha-1* was depleted in a *let-767(s2176)* loss-of-function mutant balanced with the *sDp3* duplication (Zhang et al., 2012). The single depletion of *vha-1* and *aps-1* induced a 56% and 50% increase of ERM-1 Bl/cytoplasmic ratio, respectively, and the co-depletion of these factors led to a 109% increase of this ratio, suggesting an additive effect (Fig 5A). Conversely, while the Bl/cyto ratio of ERM-1 in the balanced *let-767(s2176)* mutant was not different from control worms (ratio of 1.6 vs 1.5, respectively), this ratio was dramatically increased upon *vha-1(RNAi)* co-depletion compared to control (ratio of 4.1; 2.9-fold increase) or *vha-1(RNAi)* alone (ratio of 2.3; 1.7-fold increase) (Fig 5B-C). These results indicated that V0-ATPase genetically interacts with GSLs, but not with AP-1 in intestinal polarity maintenance.

**Figure 5.**
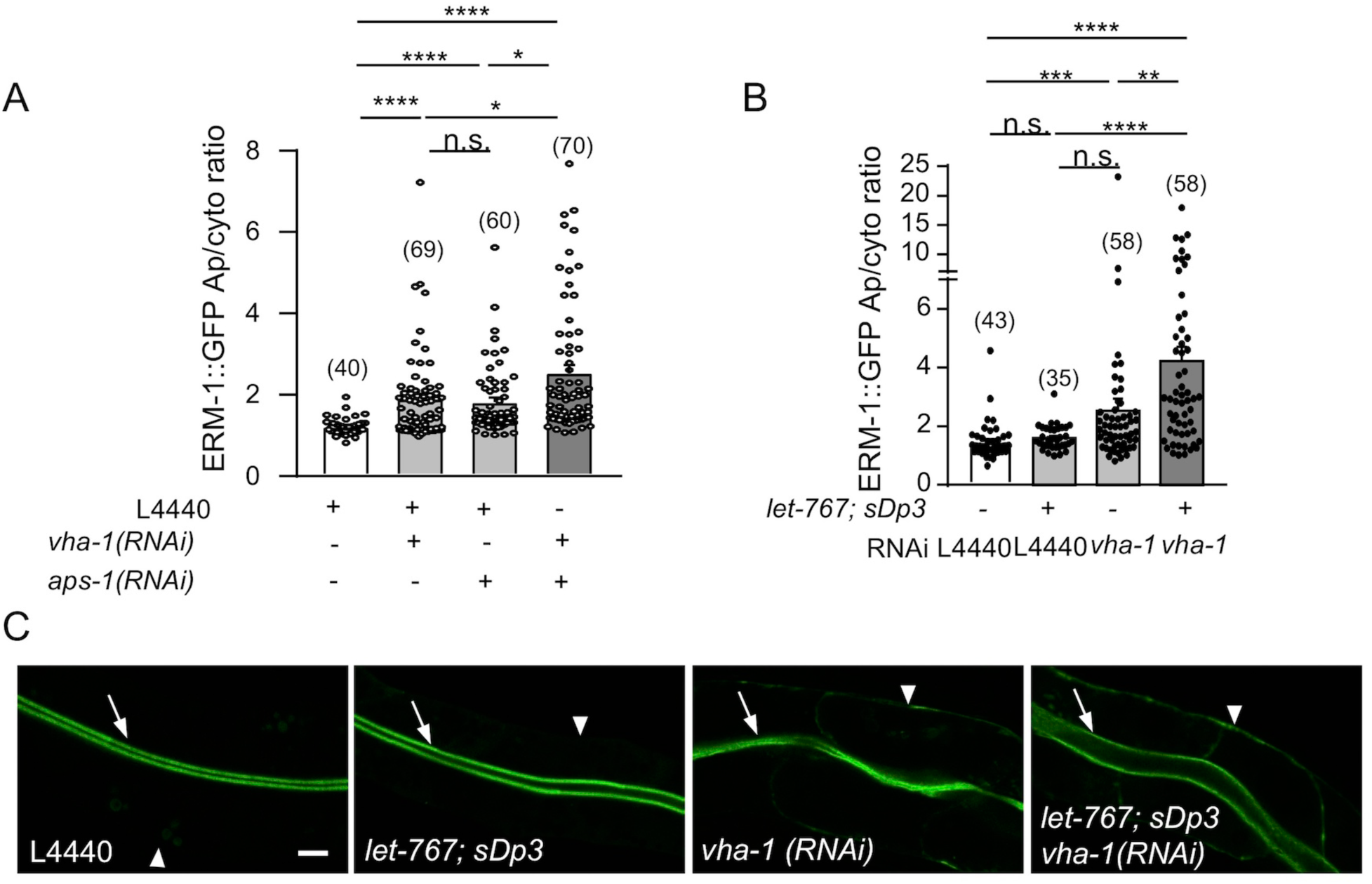
V0-ATPase genetically interacts with GSLs in intestinal polarity maintenance. (A) *vha-1* and *aps-1(RNAi)* additionally induce a basolateral mislocalization of ERM-1. *C*. *elegans* worms expressing ERM-1::GFP were silenced for control (L4440) or a mix of control, *vha-1* or *aps-1(RNAi)* as indicated and the basolateral/cytoplasmic ratio of ERM-1::GFP (Bl/cyto) was quantified. (B-C) V0-ATPase and GSLs synergistically control intestinal polarity maintenance. ERM-1::GFP expressing wild type and *let-767, sDp3* loss-of-function strains were silenced for *vha-1*. (B) shows the quantification of ERM-1::GFP Bl/cyto ratio and (C) a representative experiment. Histograms are mean ± SEM on each panel, dots represent individual worms and the total number of worms from 3 independent experiments is indicated in brackets. Arrows and arrowheads show the apical and basolateral PM, respectively. Scale bar, 5µm. Krustal-Wallis test. n.s. non-significant, *p<0.05, **p<0.01, ***p<0.001, ****p<0.0001.

### V0-ATPase depletion phenocopies Microvillus inclusion disease

Next, we asked whether the polarity defects associated with V0-ATPase depletion were associated with brush border structural defects. Combined TEM and super-resolution imaging demonstrated that prolonged V0-ATPase depletion (72 and 96h of *vha-1(RNAi)*) specifically induced a microvillus atrophy, with smaller, thinner or even absent microvilli at the apical PM of enterocytes (Fig 6A-C). Conversely, V1-ATPase depletion (*vha-8(RNAi)*), despite inducing a decreased worm size and a mild decrease of microvilli length, did not affect the microvilli structure and organization (Figs 6A-C and S3A). The severe apical PM defect induced by V0-ATPase depletion was observed along the whole surface of the deformed lumen and was often associated with a local unhooking of microvilli membrane from the underlying cytoskeleton (Fig S3B) as well as slightly lengthened adherens junctions (CeAJs) (Fig S3C-D). Thus, V0-ATPase depletion specifically leads to severe apical PM structural defects, which may affect the absorptive function of the enterocytes, in line with the luminal accumulation of cellular or bacterial debris and the absence of yolk storage granules observed in *vha-1(RNA)* but not *vha-8(RNAi)* worms (Fig S3E-F). In addition to these apical PM defects, super-resolution microscopy showed that the brush border components ACT-5 and ERM-1 adopt a pattern reminiscent of microvilli when mislocalized to the basolateral PM (Fig 6D-E) although TEM analysis did not reveal the presence of *bona fide* microvilli.

**Figure 6.**
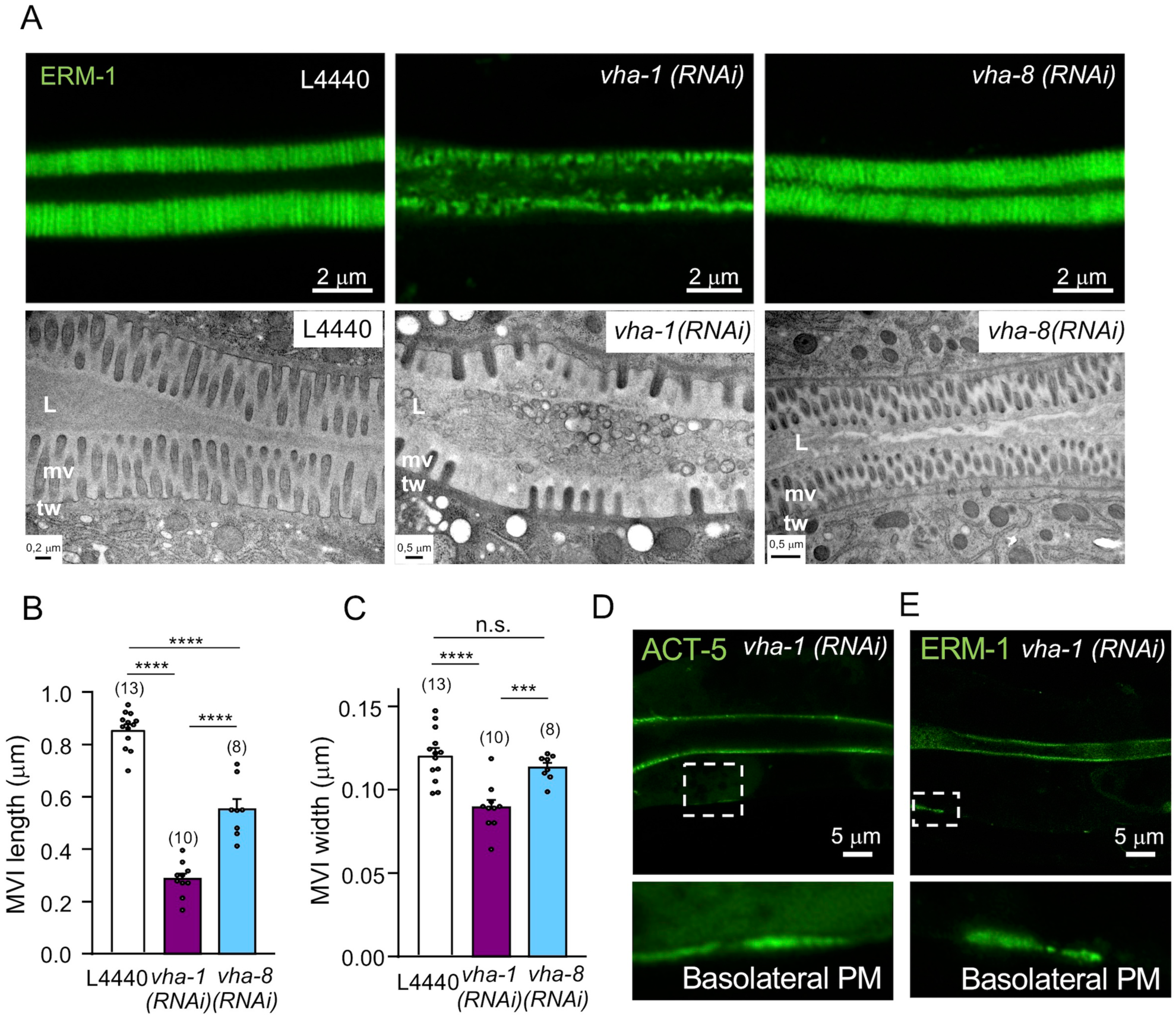
V0-ATPase depletion induces microvillus atrophy in *C*. *elegans*. (A) Visualization of microvilli by super-resolution (upper panels, 96h RNAi, ERM-1::GFP) and TEM (lower panels, 72h RNAi) analysis. Control animals showed the typical intestinal brush border morphology consisting of a dense brush border of microvilli surrounded by a glycocalyx and stabilized by the terminal web, a network of actin and intermediate filaments running below and in parallel to the apical plasma membrane. Conversely, *vha-1(RNAi)* worms display a microvillus atrophy, with irregular, shorter and sparse microvilli, a phenotype that is not observed in *vha-8* depleted worms. The terminal web appears normal in all cases. L, lumen; mv, microvilli; tw, terminal web. (B-C) Measurement of microvilli length and width (C) upon 72h V0 and V1-ATPase knockdown from TEM images. Data are mean ± SEM, n=5 microvilli/worm on the number of worms indicated in brackets. n.s., non-significant, ***p<0,001, ****p<0.0001. (D-E) The brush border components ACT-5 and ERM-1 localize to the basolateral PM with a microvilli-like pattern upon 96h *vha-1(RNAi)*. Lower panels are magnified images of the selected areas. Scale bar size is shown on each panel.

Surprisingly, prolonged V0-ATPase depletion led to the accumulation of rounded ERM-1^+^ and ACT-5^+^ structures (Fig 7A-B), which displayed a microvilli-like pattern on higher magnification (Fig 7C) and closely resemble cytoplasmic microvillus inclusions (MVIs) observed in MVID patients (Iancu et al., 2007) and in *STXBP2* KO organoids MVID model (Mosa et al., 2018). These structures are also enriched in PKC-3 and SNAP-29 (Fig 7D-E). TEM experiments performed on *vha-1* depleted worms confirmed the presence of typical MVIs lined by inward-pointing microvilli (Fig 7F). We also observed many vacuole-like structures with a wide size (0.1–0.6 μm in diameter), shape and electron-density heterogeneity and containing cellular material (Fig 7G). These vacuole-like structures were positive for the microvilli-specific activated phospho-form of ezrin, as observed by immuno-EM (Fig 7H) and are similar to the already described “irregular” MVIs that are probably immature or degrading MVIs (Iancu et al., 2007, Mosa et al., 2018). Strikingly, microvillus atrophy and MVIs are cellular hallmarks of Microvillus inclusion disease (MVID, OMIM: 251850), a congenital disorder of intestinal epithelial cells (Ruemmele et al., 2006) caused by mutations in the genes coding for the trafficking proteins Myosin 5B (MYO5B) (Muller et al., 2008), Syntaxin3 (STX3) (Wiegerinck et al., 2014) or Munc18-2 (STXBP2) (Vogel et al., 2017b). Interestingly, enterocytes of patients with MVID also displayed similar polarity defects than that observed upon *vha-1(RNAi)*, such as the basolateral appearance of microvilli-specific proteins (Wiegerinck et al., 2014), the basolateral mislocalization of PAR polarity module components (Michaux et al., 2016) as well as the cytoplasmic displacement of trafficking (i.e. STX3, the SNAREs family component SNAP23 and cellubrevin) (Dhekne et al., 2018) proteins. Finally, MVID has also been associated with enlarged lysosomes displaying a heterogeneous content (Fig 2A, C) and with the accumulation of cytoplasmic translucent vesicles (Fig 2D) (Iancu et al., 2007, Vogel et al., 2016) as well as the specific disorganization of Rab11^+^ apical recycling endosomes (Fig 2G-H) (Vogel et al., 2017a, Talmon et al., 2012). Together with the fact that *snap-29(RNAi)* also induces polarity defects (Fig 3D-G) and the formation of MVIs (Fig 7I), these observations strongly suggest that V0-ATPase depletion in *C*. *elegans* fully recapitulates the defects observed in the enterocytes of patients suffering from MVID and MVID *in vitro* mammalian models (Table S2, S3) (Mosa et al., 2018).

**Figure 7.**
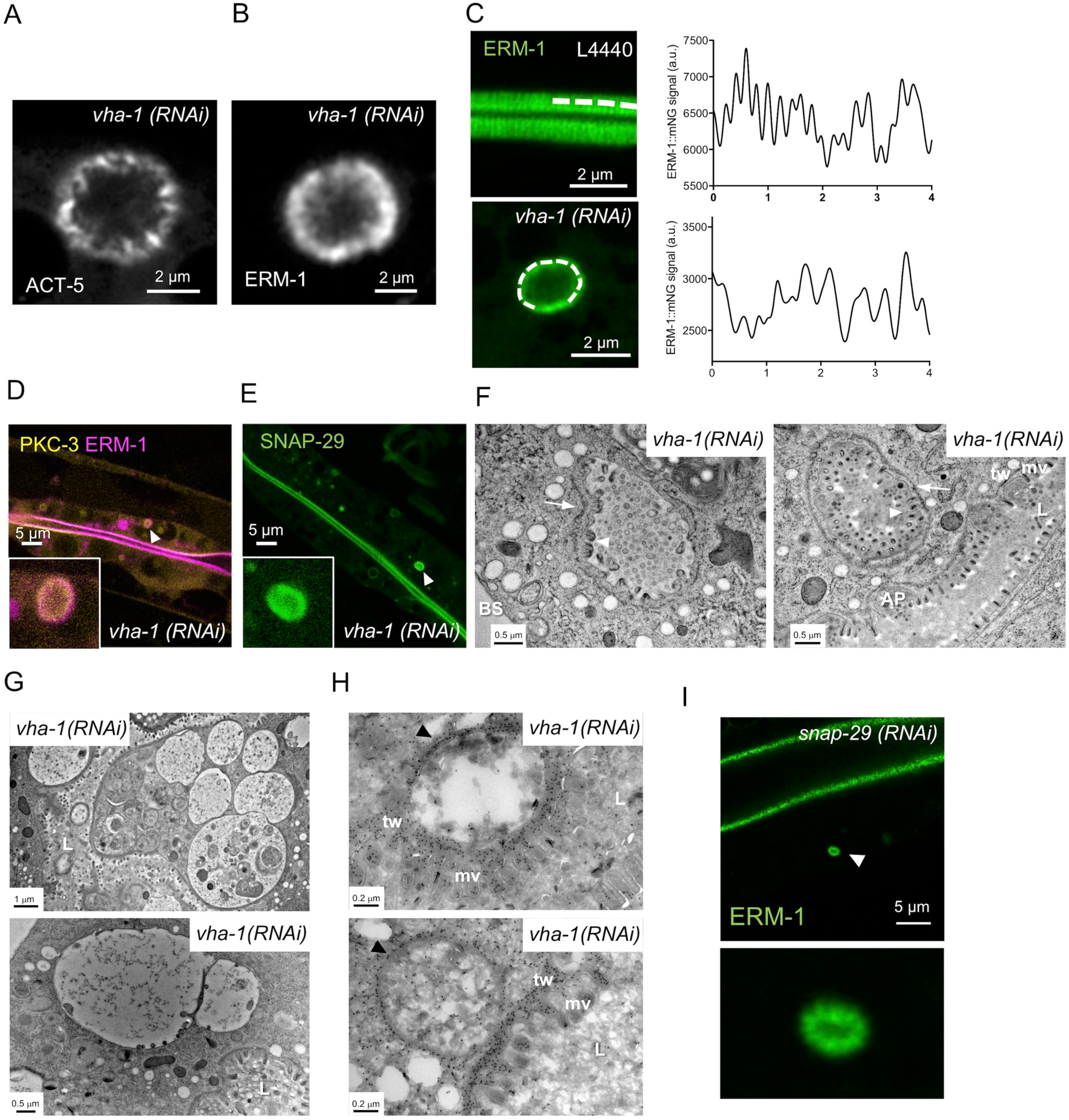
V0-ATPase silencing induces MVID in *C*. *elegans*. (A-B) 96h V0-ATPase knockdown (*vha-1(RNAi)*) induces the formation of cytoplasmic ERM-1^+^ and ACT-5^+^ microvillus inclusion-like structures. (C) Line scan shows that rounded structures found in the cytoplasm (lower panel) display periodic pattern reminiscent of apical microvilli (upper panel). A line scan of 4 μm was performed (dotted line) on ERM-1::mNG expressing worms apical PM (control worms) and on ERM-1::mNG^+^ MVI-like structures (96h of *vha-1(RNAi)*). (D-E) GFP::PKC-3 and SNAP-29::GFP localize to *vha-1(RNAi)* induced MVI-like structures. Insert is a magnified image of the structure indicated by the arrowhead. (F-H) V0-ATPase silencing mostly induces cytoplasmic ERM-1^+^ vacuoles/irregular MVIs as well as typical MVIs. (F-G) Ultrastructural analysis of cytoplasmic inclusions observed in *C*. *elegans* silenced for *vha-1* during 96h by TEM. Note the classical MVIs showing the symmetrical, mainly inward-pointing arrangement of microvilli (arrowheads) and the area around the inclusion corresponding to the terminal web (arrows). Typical MVIs were found in 3/16 worms. (G) Numerous and larger ‘irregular’ vacuole-inclusions with a wide heterogeneity in size, shape, and electron-density were found upon 96h *vha-1* silencing. The lumen contains probably amorphous, microvesicular, membranous and lipid components. (H) Immuno-EM analysis of vacuole-like structures using phospho-ezrin antibody and nanogold protein A gold in *vha-1(RNAi)* worms (96h) (dark dots). Anti-phospho-ezrin antibody accumulates around *vha-1(RNAi)*-induced vacuole-like structures and on the apical PM. (I) *snap-29(RNAi)* (96h) also induced the formation of cytoplasmic ERM-1::mNG^+^ MVIs. Lower panel is a magnified picture of the MVI indicated by the arrowhead. L, lumen; mv, microvilli; tw, terminal web; AP, apical PM; BS, basal PM. Scale bar is shown on each panel.

### Study of MVIs dynamics in *C*. *elegans* model

Using live microscopy in intestinal organoids derived from *STXBP2* knock-out mouse, we recently contributed to show that MVIs can arise *de novo* in the cytoplasm or by internalization of both the apical and the basolateral PM (Mosa et al., 2018). To test whether these various modes of MVI onset are conserved in *C*. *elegans* upon V0-ATPase depletion, we performed live imaging on endogenously expressed ERM-1::mNG after 96h of *vha-1(RNAi)*. This experiment confirmed that MVI-like structures can rapidly nucleate in the cytoplasm (∼3 min) and remained in the cytoplasm from less than 3 min to more than 1h (Fig 8A and Movie S1-S2), a process that may involve lysosomal remodelling (Mosa et al., 2018). These structures may also arise by internalization of large ERM-1::GFP^+^structures from the apical and basolateral PM of *vha-1* depleted worms (Fig 8B, S4 and Movie S3), which has been linked to an autophagocytic process (Reinshagen et al., 2002). Since autophagosomes and lysosomes have been linked to MVIs, we hypothesized that these structures undergo a cytoplasmic elimination through these degradation pathways. Consistently, live tracking of MVIs in *STXBP2* KO organoids revealed that MVIs are often surrounded by LAMP2^+^ membranes and mostly disappear by cytoplasmic desintegration while the rest was eliminated by PM fusion or cell shedding (Mosa et al., 2018). Interestingly, we found that irregular MVIs induced by *vha-1(RNAi)* were often in close proximity to mixed organelles/lysosomes (Fig 8C) and that some of the ERM-1^+^ MVIs were also positive for the lysosomal marker LMP-1 (Fig 8D). Furthermore, some MVIs were also surrounded by a second membranous layer positive for ERM-1 (Fig 8E). Considering that i) ezrin has been shown to accumulate at the phagosome membrane and regulate phago-lysosomal fusion (Marion et al., 2011) and ii) V0-ATPase extinction dramatically activates autophagy, as ascertained by the relocalization of LGG-1::GFP (LC3 *C*. *elegans* ortholog) from the apical PM to cytoplasmic punctaes (Fig 8 G-H) (Chang et al., 2017), these data strongly suggest that MVIs are eliminated by autophagosomes and/or lysosomes in *C*. *elegans* intestine, as proposed in mammals. This was confirmed by timelapse microscopy on ERM-1^+^ MVIs, which progressively disintegrated in the cytoplasm or inside ERM-1^+^ secondary structures (Fig 8F and Movie S4).

**Figure 8.**
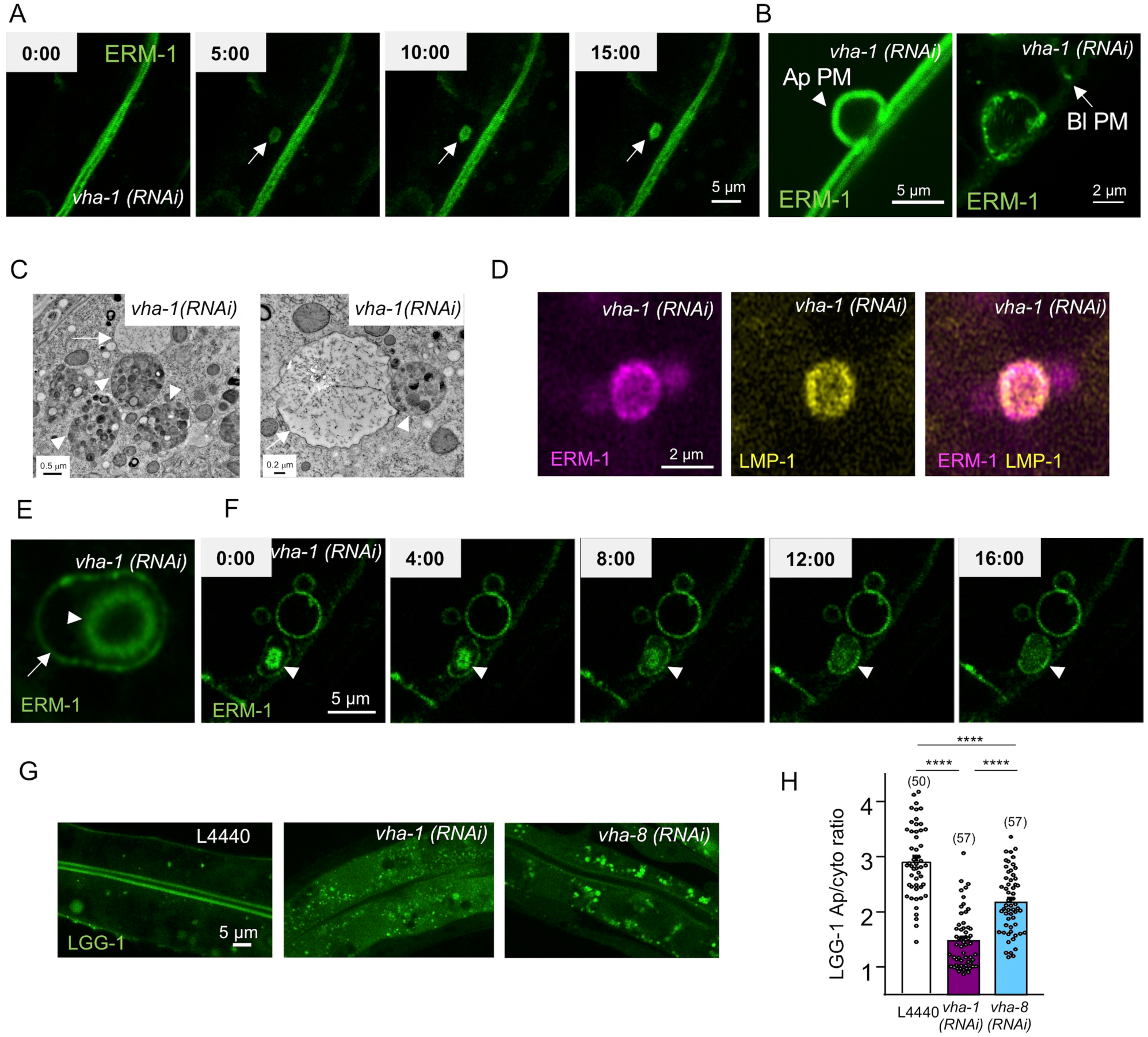
Dynamics of microvillus inclusions in *C*. *elegans*. (A) MVIs cytoplasmic nucleation observed by timelapse imaging in endogenous ERM-1::mNG expressing worms silenced for *vha-1* during 96h. Snapshots are shown every 5 min, MVI is indicated by the arrow. (B) MVIs can arise by internalization of the apical or basolateral PM. ERM-1::GFP expressing worms were silenced for *vha-1* during 72h and imaged by conventional (left panel) or super-resolution (right panel) microscopy. Arrowhead and arrow show the apical and basolateral PM, respectively. (C) Irregular MVIs/vacuoles (arrow) are often in the vicinity of lysosomes/mixed-organelles (arrowhead). N2 worms were silenced for *vha-1* during 96h and analyzed by TEM. (D) ERM-1::wSc and LMP-1::GFP colocalize at *vha-1(RNAi)*-induced MVIs (96h). (E-F) 96h *vha-1(RNAi)*-induced MVIs can be surrounded by a second ERM-1^+^ layer in which MVIs are degraded. (E) shows a super-resolution image of an MVI (arrowhead) and the surrounding ERM-1^+^ layer (arrow), (F) shows a timelapse imaging of *vha-1(RNAi)*-induced MVI disappearance (image every 4 min). (G-H) Autophagy is activated more upon *vha-1(RNAi)* than *vha-8(RNAi)*. LGG-1::GFP expressing worms were silenced for *vha-1* or *vha-8* during 72h and imaged. (H) shows the quantification of LGG-1::GFP apical/cytoplasmic (Ap/cyto) ratio. Histograms are mean ± SEM on each panel, dots represent individual worms and the total number of worms from 3 independent experiments is indicated in brackets. ****p<0.0001.

## Discussion

By performing a double-RNAi screen in *C*. *elegans* intestine model we identified a new function of the V0-ATPase complex as a critical determinant for the maintenance of one of the most prominent cell surfaces, the absorptive apical PM of enterocytes. Our results demonstrate that the V0-ATPase ensures the apical localization of polarity modules, brush border components and trafficking factors, a major role that is substantiated by the severe brush border defects consecutive to V0-ATPase depletion.

This study confirms the biological significance of systematic RNAi screens in *C*. *elegans* to understand the complex mechanisms of polarity maintenance. Indeed, this model has allowed to characterize the major role of membrane trafficking factors in epithelial polarity maintenance including GSLs (Zhang et al., 2011), clathrin and its adaptor AP-1 (Shafaq-Zadah et al., 2012, Zhang et al., 2012), the kinase Lats/WTS-1 (Kang et al., 2009) and the interactions between polarity proteins and membrane trafficking (Winter et al., 2012, Balklava et al., 2007). Intriguingly, while interfering with Clathrin/AP-1, PAR-5 or GSLs pathways induced a RAB-11^+^ endosomes loss and an inversion of both apical and basolateral PM proteins polarity (Shafaq-Zadah et al., 2012, Zhang et al., 2012, Zhang et al., 2011, Winter et al., 2012), we showed here that V0-ATPase silencing has a specific effect on brush border and CDC-42/PAR polarity but not on transmembrane or basolateral proteins (Figs 2, 3). This surprising result points to a specific role of V0-ATPase in late trafficking events, downstream of sorting factors such as GSLs, clathrin and AP-1, which is consistent with the deleterious effect of GSLs biosynthesis enzyme or *aps-1* depletion on the localization of the V0-ATPase subunit VHA-6 apical localization (Zhu et al., 2015). This conclusion is supported by i) the known function of V0-ATPase in vesicle fusion to the plasma membrane and its localisation in microvilli, ii) the accumulation of vesicles upon V0-ATPase depletion, iii) the role of the SNARE SNAP-29 downstream of the V0-ATPase and iv) the phenotypic similarities with the loss of Myosin 5B, STX3 or Munc18-2 in patients; indeed all these factors have been found to control late trafficking events just upstream of, or during, the fusion to the apical PM. In summary, clathrin/AP-1 (Shafaq-Zadah et al., 2012, Zhang et al., 2012), GSLs (Zhang et al., 2011, Zhang et al., 2012) or the kinase Lats/WTS-1 (Kang et al., 2009) might control sorting events, while V0-ATPase, Myosin 5B, STX3 or Munc18-2 would control late trafficking steps (Fig S5).

Furthermore, the genetic interaction experiments performed in this study advocate that the two known sorting pathways, AP-1 and GSLs, converge towards RAB-11^+^ endosomes and the V0-ATPase to ensure the polarized localization of soluble factors necessary for intestinal polarity maintenance (i.e. the PAR module and ERM-1). First, AP-1 could regulate an early sorting step, which may rely on the recognition of sorting signals in cargo proteins sequence, as shown previously for AP-1-dependent basolateral sorting through di-leucin or tyrosine-based motifs (Bonifacino, 2014); the additional effect following the loss of AP-1 and V0-ATPase suggests that the latter controls more than just AP-1-dependent cargos. Second, GSLs could control a parallel early sorting step based on other signals (i.e. GPI anchor) in a pathway encompassing also a V0-ATPase-dependent apical PM exocytosis, based on the synthetic effect observed. Hence, we propose that polarity module and brush border components undergo a complex apical PM targeting mechanism at both the TGN and along the secretory/recycling pathways.

The precise mechanism by which the V0-ATPase controls the maintenance of the intestinal polarity remains to be elucidated. The most studied function of this complex in membrane trafficking relies on the H^+^-pump-dependent (V0+V1 subunits) acidification of intracellular organelles, which ensures their maturation and transport capacity (Maxson and Grinstein, 2014). However, our results demonstrated a specific requirement for its V0-sector in polarity maintenance, in an acidification-independent manner and likely through its already characterized function in vesicle exocytosis (Hiesinger et al., 2005, Di Giovanni et al., 2010, Baars et al., 2007, Liegeois et al., 2006). Indeed, the functional relevance of V0-ATPase interaction with the fusion machinery has been demonstrated in many species, such as drosophila or *T*. *marmorata*, where the V0-a1 subunit (V100 in drosophila) has been shown to interact with the t-SNAREs Syx and SNAP25 (Hiesinger et al., 2005) and with the v-SNARE VAMP-2 (Morel et al., 2003), respectively. Furthermore, the direct interaction between the V0-c subunit (the ortholog of *vha-1*) and VAMP2 has been shown to be indispensable for efficient synaptic vesicle fusion in mammalian cells (Di Giovanni et al., 2010). A similar mechanism could take place for the apical exocytosis of vesicles containing either polarity and brush border components, which may directly interact with the V0-ATPase, and/or apical adaptor proteins, such as the ezrin partner NHERF-1/EBP50 that has been recently found associated with the V-ATPase in interactome studies (Merkulova et al., 2015). In line, we found that *vha-1(RNAi)* decreased the apical confinement of the SNARE SNAP-29, a member of the SNAP23/25/29 family (Steegmaier et al., 1998) which seems to be mislocalized in MVID mammalian models (Dhekne et al., 2018), and that *snap-29* extinction phenocopies that of *vha-1* on *C*. *elegans* intestinal polarity. Considering that SNAP-29 has been localized to RAB-11^+^ endosomes in *C*. *elegans* (Kang et al., 2011), likely at the terminal web (Fig 3) and that mammalian SNAP29 physically interacts with STX3 (Steegmaier et al., 1998), the V0-ATPase could directly interact with SNAP-29 to target the exocytosis of RAB-11^+^ endosomes and be involved in the same pathway than that of Myosin 5B/STX3/Munc18-2, which may also involve other trafficking factors, such as the N-ethylmaleimide sensitive factor NSF-1/NSF (S1 Table).

Beyond disturbing polarity, we uncovered that V0-ATPase depletion induced severe microvilli defects and the formation of MVIs that are typical features of MVID. MVID is a devastating orphan congenital disorder of intestinal cells that manifested in intractable diarrhoea and intestinal absorption defects appearing in early life of new-borns (Halac et al., 2011, Ruemmele et al., 2006) and therapeutic avenues presently only rely on total parenteral nutrition or small bowel transplantation (Halac et al., 2011). MVID originates from mutations of MYO5B, STX3 or STXBP2 (Dhekne et al., 2018) that belong to the same apical trafficking pathway. At the molecular level, a defective apical PM tethering of RAB-8^+^/RAB-11^+^ endosomes by the molecular motor Myosin 5B (Knowles et al., 2014) which interacts with STX3 (Vogel et al., 2015) in concert with the sec/munc family protein Munc18-2 that plays an accessory role for STX3-dependent vesicle tethering Munc18-2 (Vogel et al., 2017b) has been proposed to be the major pathway leading to MVID phenotype. Hence, MVID is strictly linked to a polarized trafficking defect and *in vitro* mammalian 2D or 3D cell culture models have been developed to study its ill-defined underlying mechanisms, based on Myosin 5B or Rab8/Rab11 depletion and disruption of Myosin 5B-Rab8/11 interaction (Feng et al., 2017, Knowles et al., 2014, Engevik et al., 2018, Kravtsov et al., 2016, Weis et al., 2016, Vogel et al., 2015, Schneeberger et al., 2015) as well as STX3 (Wiegerinck et al., 2014, Vogel et al., 2015) or STXBP2 (Mosa et al., 2018, Vogel et al., 2017b) mutations.

However, some discrepancies were found while trying to integrate polarity maintenance mechanisms identified *in vitro* and *in vivo*. For instance, while AP-1 was renowned for basolateral sorting in cultured cells (Gravotta et al., 2012), its critical role in apical traffic was characterized *in vivo* in *C*. *elegans* and in mouse (Shafaq-Zadah et al., 2012, Zhang et al., 2012, Hase et al., 2013). While the *in vitro* culture of intestinal organoids derived from MVID patients or MVID causing genes-deficient mouse (Schneeberger et al., 2018, Mosa et al., 2018, Wiegerinck et al., 2014) and the mutation of *myosin5b*/*goosepimples* in zebrafish (Sidhaye et al., 2016) seem to reproduce most of the cellular and structural hallmarks of MVID, these models are not suitable for systematic studies. Conversely, the *in vivo*, easy-to-use and screening amenable *C*. *elegans* model has been widely used to understand intestinal polarity mechanisms (Zhu et al., 2015, Zhang et al., 2012, Winter et al., 2012, Shafaq-Zadah et al., 2012, Kiela and Ghishan, 2016, Sobota et al., 2009, Kang et al., 2009), but single deletion of the orthologs of MVID-causing genes (i.e. *hum-2*/*MYO5B, unc-64*/*STX3*) seems not to induce an MVID phenotype (our unpublished observations), probably due to compensatory mechanisms. We describe here for the first time a *C*. *elegans* model, V0-ATPase depletion, that fully phenocopies the hallmarks of MVID (Table S3) and that could be used to better characterize polarity maintenance mechanisms and MVID pathophysiology. As a proof of principle, we studied the dynamics of MVIs, which onset and elimination are still enigmatic. According to the “combined” model of MVI onset (Schneeberger et al., 2018), in a recent paper using *STXBP2* KO organoids (Mosa et al., 2018) and in the present study (Fig 7), we showed that MVIs may arise by two main mechanisms; *de novo* nucleation in the cytoplasm or internalization of the PM. Both may rely on the failure of apical exocytosis from RAB-11^+^ endosomes. Indeed, this defect would on one hand result in the sequestration of polarity factors in the cytoplasm, that may be an upstream signal to target newly-synthesized apical PAR, PM proteins and cytoskeleton and consequently nucleate a new apical PM in the cytoplasm on accumulating vesicles. On the other hand, defective exocytosis may lead to a local apical PM protein/lipid homeostasis defect that would create a signal for endocytosis and targeting to autophagosomal-lysosomal degradation, as proposed before (Reinshagen et al., 2002) and verified in this report (Fig 8). Finally, we provide compelling evidence that V0-ATPase depletion fully phenocopies MVID and this suggests that V0-ATPase may be involved in the aetiology of MVID and other absorption disorders associated with apical PM defects, or affected in patients (Table S3). Complementary use of both *C*. *elegans*, as a tool to screen for correctors of polarity maintenance defects, as well as mammalian intestinal organoids or human samples to confirm the conservation of V0-ATPase function, may lead to a better understanding of the mechanisms of polarity maintenance as well as that of rare genetic enteropathies.

## Materials and methods

### *C*. *elegans* strains and maintenance

Strains were maintained under typical conditions as described (Brenner, 1974). mNG and mScarlet-tagged ERM-1, UNC-64 and PGP-1 were generated at the « Biologie de *Cænorhabditis elegans* » facility (UMS3421, Lyon, France). The strains used in this study are listed in Table S4.

### Gene silencing and screening by RNAi

Larval RNAi was performed by feeding as described using the Ahringer-Source BioScience library (Kamath and Ahringer, 2003, Shafaq-Zadah et al., 2012). Briefly, adult hermaphrodites were bleached and transferred to RNAi plates containing HT115 *E*. *coli* bacteria expressing the L4440 plasmid alone (control) or encoding the RNAi. For co-depletion of *vha-1* and *aps-1*, equal amount of exponential phase bacterial culture (OD_600_ ∼0.6) were mixed together or with L4440 (single depletion). Developing worms were kept in RNAi for 72-96h before mounting and imaging. RNAi specificity was verified by sequencing and efficiency by target::GFP signal disappearance, as shown in Figures.

### *in vivo* confocal imaging using *C*. *elegans*

*C*. *elegans* imaging was performed *in vivo* by mounting the worms on a 10% agarose pad in a solution of 100 nm polystyrene microbeads (Polysciences Inc.) to stop worm movement. Confocal observations were performed using a Leica (Wetzlar, Germany) SPE equipped with a 63X, 1.4 NA objective (LAS AF software) or a super-resolution Zeiss (Oberkochen, Germany) LSM880 + Airyscan equipped with a 63X, 1.4 NA objective (Zen Black software). All images were examined using Fiji software.

### TEM

Control and RNAi worms (72-96 hours) were fixed by high-pressure freezing with EMPACT-2 (Leica Microsystems) and then freeze substituted in anhydrous acetone containing 1% OsO4, 0.5% glutaraldehyde and 0.25% uranyl acetate for 60 h in a freeze substitution system (AFS-2; Leica Microsystems). Larvae were embedded in Epon-Araldite mix (EMS hard formula). Adhesive frames were used (11560294 GENE-FRAME 65 µL, Thermo Fisher Scientific) for flat-embedding, as previously described (Kolotuev et al., 2012), to gain better antero-posterior orientation and sectioning. Ultrathin sections were cut on an ultramicrotome (UC7; Leica Microsystems) and were collected on formvar-coated slot grids (FCF2010-CU, EMS). Each larva was sectioned in 5 different places with ≥ 10 µm between each grid to ensure that different cells were observed. Each grid contained at least 10 consecutive sections of 70 nm. TEM grids were observed using a JEM-1400 transmission electron microscope (JEOL, Tokyo, Japan) operated at 120 kV, equipped with a Gatan Orius SC 1000 camera (Gatan, Pleasanton, USA) and piloted by the Digital Micrograph program. Micrographs were analyzed using Fiji software.

### Immuno-EM

Immuno-electron microscopy was performed as described (Nicolle et al., 2015). For ultrathin cryosections and immunogold labeling, larvae were fixed with 4% paraformaldehyde plus 0.1% glutaraldehyde in M9 buffer. Worms were embedded in 12% gelatin and infused in 2.3 M sucrose. Gelatin blocks were frozen and processed for ultracryomicrotomy (UC7cryo; Leica Microsystems). Ultrathin cryosections (80 nm) were double-immunogold labeled and analyzed by TEM as described above. We applied primary antibodies: rabbit anti-phospho-ezrin (Thr567) / radixin (Thr564) / moes (Thr558) (1/100, 3149S Ozyme), detected using protein A conjugated to 10 nm gold particles (PAG, 1/50, CMC, Utrecht, the Netherlands).

### Quantification

Apical/cytoplasmic and basolateral/cytoplasmic ratios were calculated by quantifying the signal intensity on a line covering the width of the apical PM and covering at least the length of 2 adjacent intestinal cells. The intensity of the same line was quantified in the cytoplasm underneath the PM. Basolateral ERM-1 appearance was calculated using the Plot Profile function of Fiji, with a R.O.I. of 1.3 μm long crossing the highest basolateral PM intensity. The 5 highest values were kept and divided by the mean fluorescence of the same R.O.I. in the cytoplasm. Microvilli length and width were quantified using Fiji on 1) TEM pictures on at least 5 sections per worm (Fig 4) or 2) on confocal pictures by calculating the width of at least 4 lines of >100nm long/image (Fig 7). The number of RAB-5+ vesicles was quantified using a home-made plugin for Fiji, as described (Gillard et al., 2015).

### Statistical analysis

Results are presented as mean ± SEM of the number of independent experiments indicated in the legends, and scattered dots represent individual worms. The total number of larvae used is indicated in brackets in histograms. P values were calculated with the means of at least 3 independent experiments by two-tailed unpaired student’s t-test and a 95% confidence level was considered significant. Normal distribution of data and homogeneity of variance were validated by calculating the skew factor (−2 > skew < 2) and the F-test, respectively. Mann-Withney U-test and Krustal-Wallis test were used for calculating the P-values of non-normal distributions and Welch correction was applied to normal distributions with non-homogenous variances, as indicated in legends.

## Acknowledgements

We thank Verena Gobel, Ken Kemphues, Jeremy Nance, Emily Troemel, Barth Grant and Ken Sato for strains. We also thank the « Biologie de *Cænorhabditis elegans* » facility (UMS3421, Lyon, France) which generated several CRISPR/Cas9 genome-edited strains used in this study. Some strains were provided by the CGC, which is funded by NIH Office of research Infrastructure Programs (P40 OD010440; University of Minnesota, USA). We thank Aurélien Perrin for the initial PAR-6::GFP screen, Raphaël Dima and Justine Cailloce for help with screen validation as well as Anne Pacquelet for critical reading of the manuscript and helpful discussions. Imaging was performed at the photonic and electron microscopy facilities of the Microscopy Rennes imaging Center (MRiC), Rennes, France. This work was supported by the Ligue Contre le Cancer Grand Ouest [22/29/35/72/41] and the Fondation Maladies Rares. ABM is the recipient of a postdoctoral fellowship from La Ligue Nationale Contre le Cancer (2017-2018).

## Authors contribution

ABM, ON and MH performed and analyzed the experiments. ABM and GM designed the study and wrote the manuscript. GM supervised the study.

## Conflict of interest

The authors declare no conflict of interest.

## Supporting information

**Fig S1.**
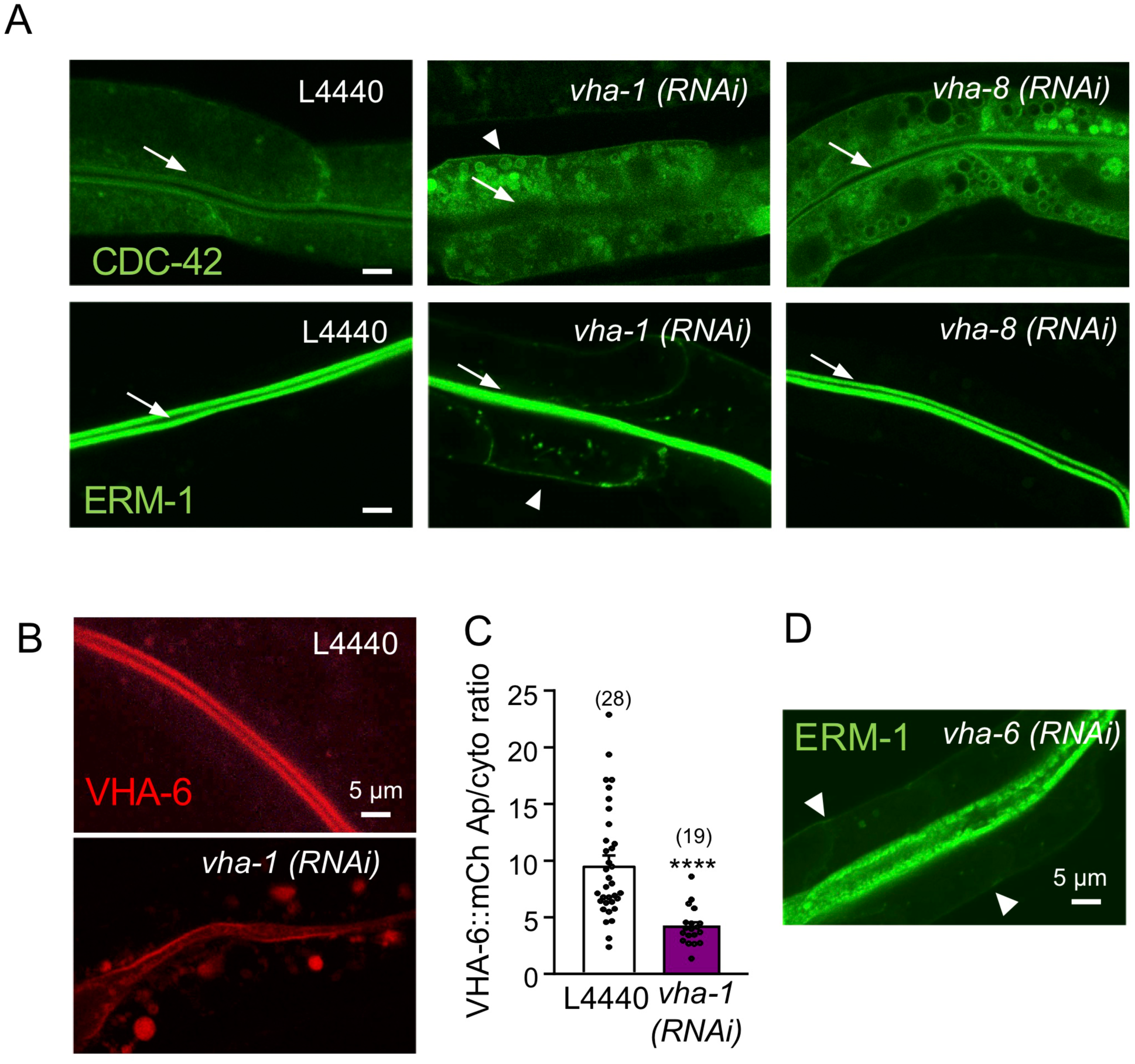
V0-ATPase silencing affects the polarized localization of polarity modules and brush border components. (A) Polarity screen results. GFP::CDC-42 (upper panels) and ERM-1::GFP (lower panels) expressing strains were silenced for control (L4440), V0-ATPase (*vha-1*) or V1-ATPase (*vha-8*) specific RNAi during 72h and imaged. Note the basolateral accumulation of both markers upon *vha-1(RNAi)*. (B-C) Effect of 72h *vha-1(RNAi)* on the expression of the V0-ATPase subunit VHA-6::mCh. (B) shows representative images and (C) the quantification of VHA-6::mCh apical/cytoplasmic ratio. The histogram shows the mean ± SEM, dots represent individual worms and the total number of worms from 3 independent experiments is indicated in brackets. ****p<0.0001. (D) 72h *vha-6(RNAi)* also induces a basolateral mislocalization of ERM-1::GFP (arrowheads). Arrows and arrowheads show the apical and basolateral PM, respectively.

**Fig S2.**
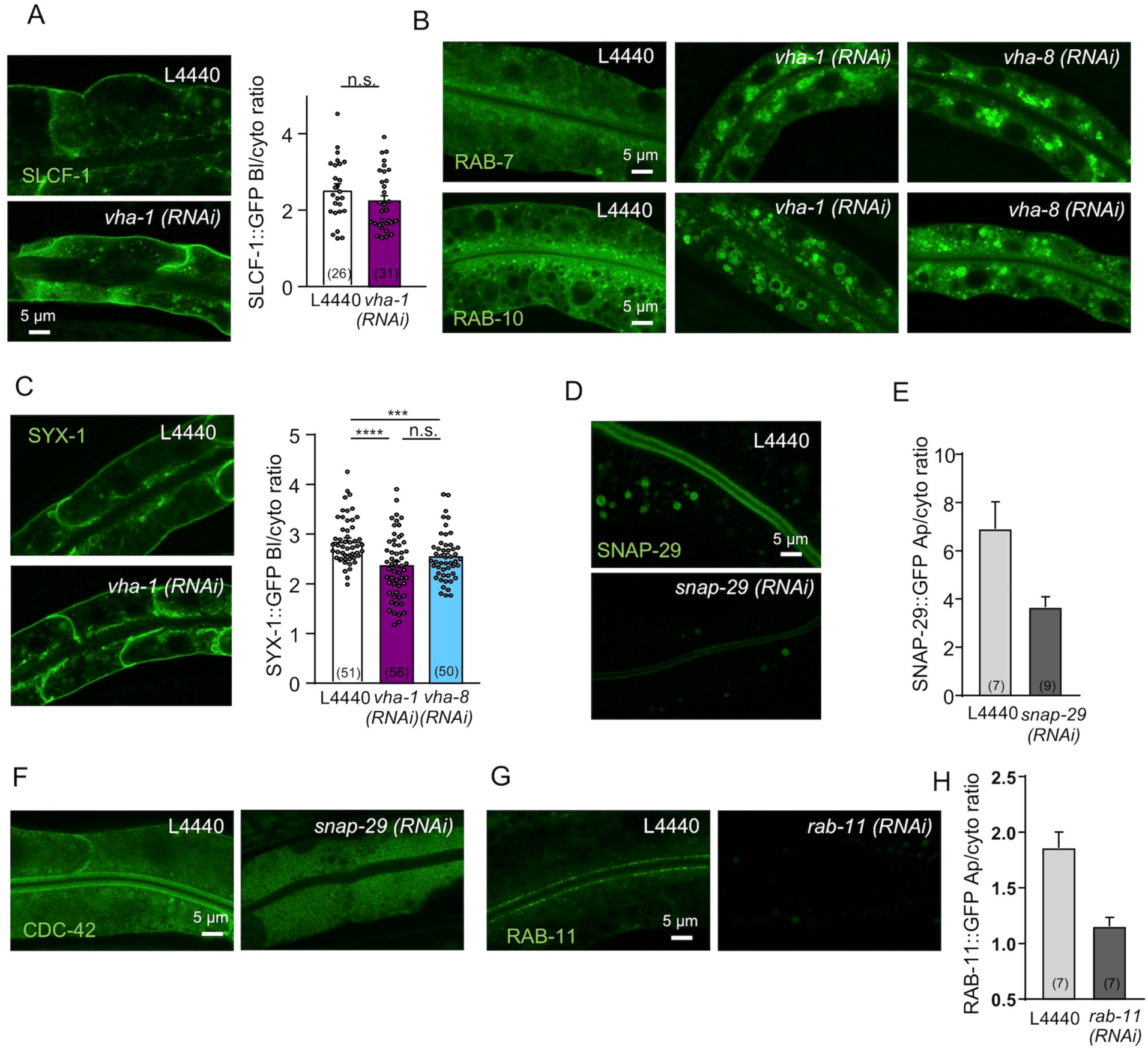
V0-ATPase polarity maintenance pathway. (A) Effect of 72h V0- or V1-ATPase silencing on the basolateral/cytoplasmic (Bl/cyto) ratio of SLCF-1. Left panel shows a representative experiment. (B) Both *vha-1* and *vha-8(RNAi)* (72h) induce an accumulation of GFP::RAB-7 and GFP::RAB-10 in large cytoplasmic structures. Image shown is representative of 52-66 RAB-7::GFP and 38-58 RAB-10::GFP expressing worms observed for each condition, out of 3-5 independent experiments. (C) Effect of 72h V0- or V1-ATPase silencing on the basolateral/cytoplasmic (Bl/cyto) ratio of SYX-1/SYN-1. Left panel shows a representative experiment. (D-F) *snap-29(RNAi)* decreases SNAP-29::GFP expression and CDC-42::GFP apical localization. (E) represents the apical/cytoplasmic ratio of SNAP-29::GFP. (G-H) *rab-11(RNAi)* decreases RAB-11 expression. (H) shows the quantification of RAB-11::GFP apical/cytoplasmic ratio. Histograms are mean ± SEM on each panel, dots represent individual worms and the total number of worms from 3 (A, C) or 1 (E, F) experiment(s) is indicated in brackets. n.s. non-significant, ***p<0,001, ****p<0,0001.

**Fig S3.**
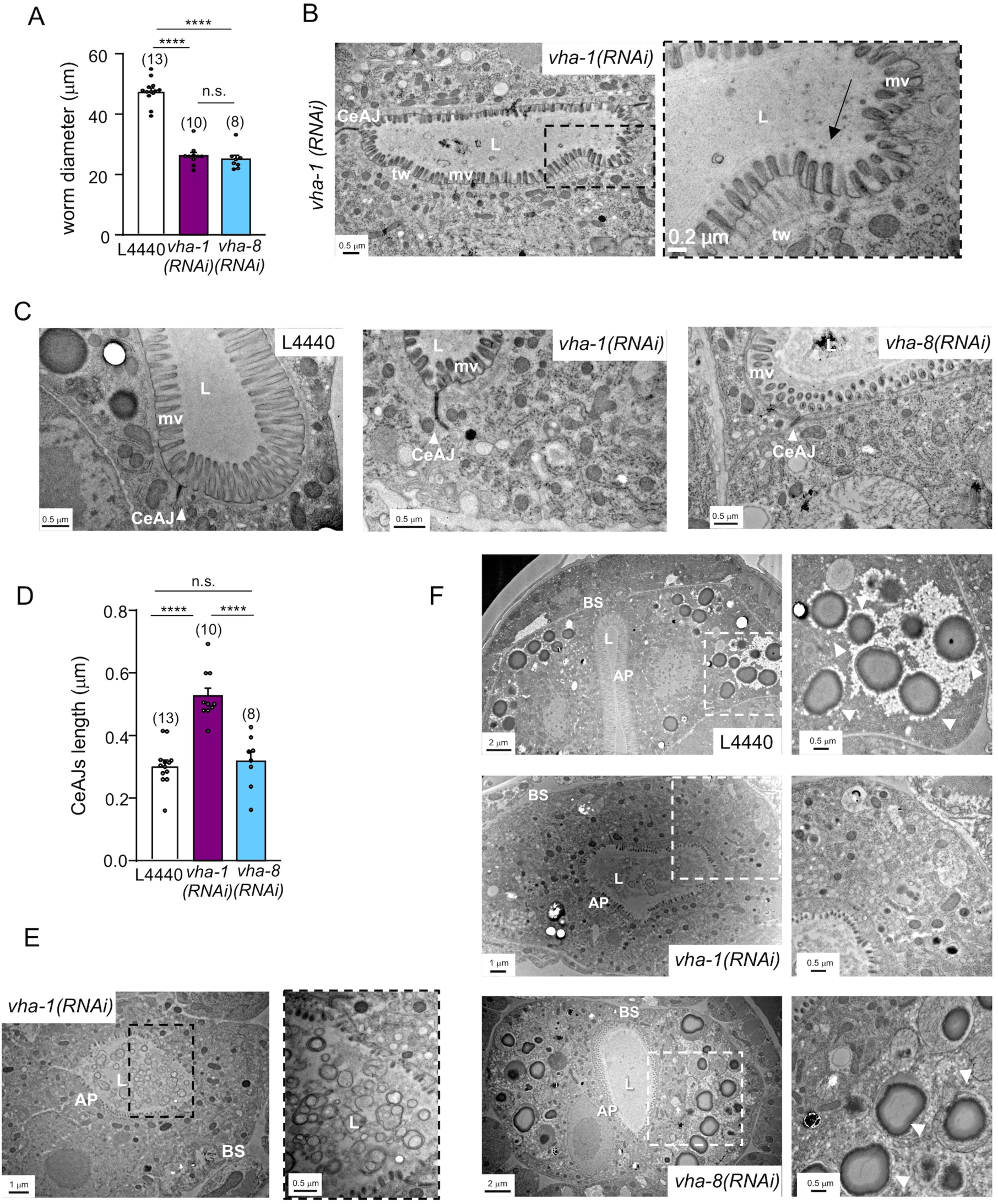
Ultrastructural analysis of V0-ATPase depletion associated defects. (A) V0- or V1-ATPase subunits silencing affects *C*. *elegans* development. Histogram shows the measurement of the worms’ diameter from TEM images. (B) *vha-1* silencing often induced a detachment of the PM from the terminal web as well as (C-D) increased the length of cell junctions (CeAJ). (D) shows the measurement of the electron-dense cell-cell junction length on TEM pictures. (E) *vha-1* depleted worms accumulate cell and/or bacterial debris in the intestinal lumen. (F) *vha-1* depleted N2 worms, contrary to control or *vha-8*-depleted worms, are devoid of yolk storage granules (arrows). L, lumen; mv, microvilli; tw, terminal web; AP, apical PM; BS, basal PM; CeAJ, *C*. *elegans* adherens junctions. Histograms show the mean ± SEM, dots represent individual worms and the total number of worms is indicated in brackets. n.s. non-significant, ****p<0,0001.

**Fig S4.**
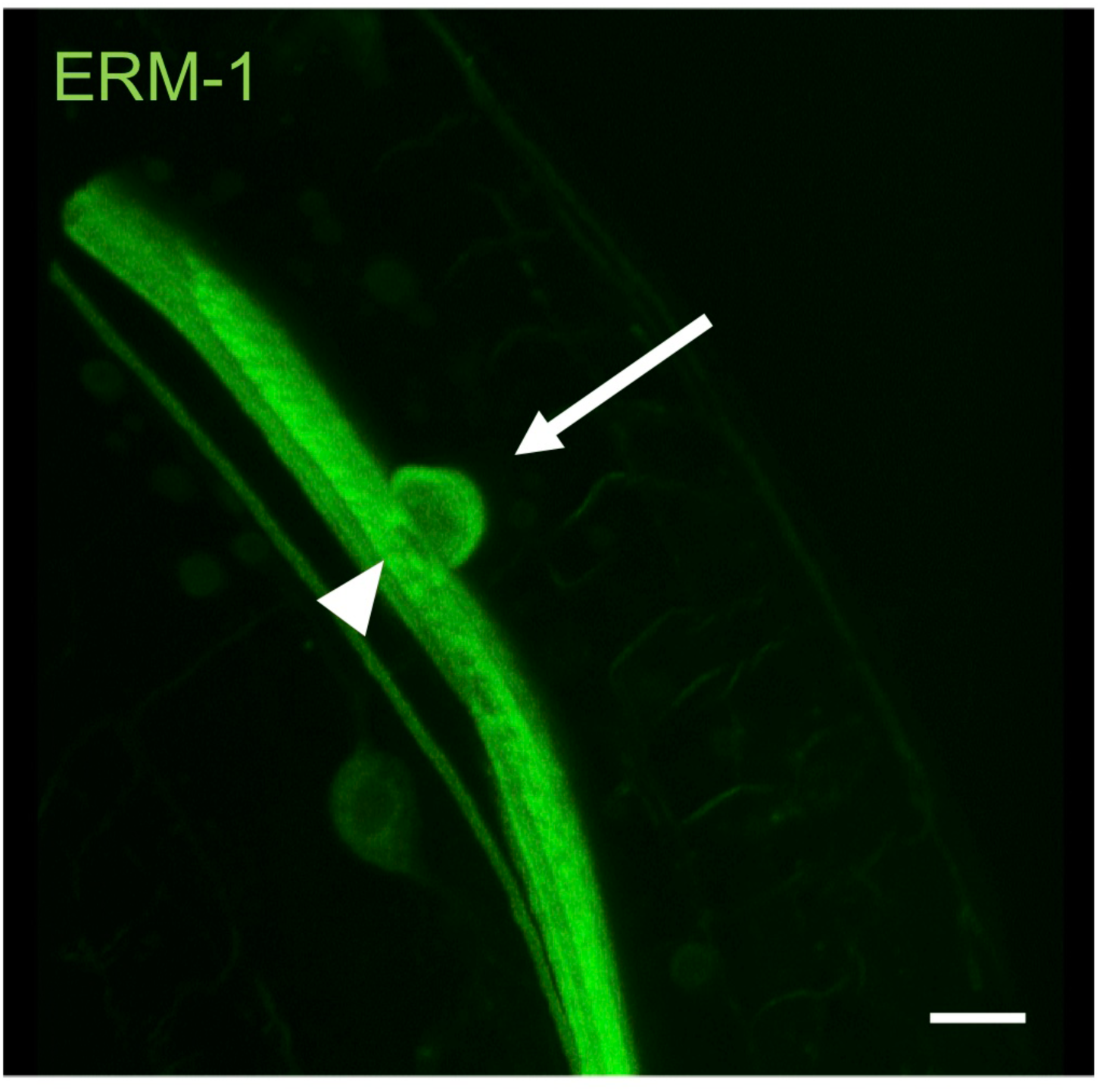
MVIs onset by internalization of the apical PM. 3D projection of ERM-1::GFP expressing worms silenced for *vha-1* during 72h. The arrow indicates the internalization of the apical PM. Note the large hole induced by the PM invagination (arrowhead). Scale bar: 5 μm.

**Fig S5.**
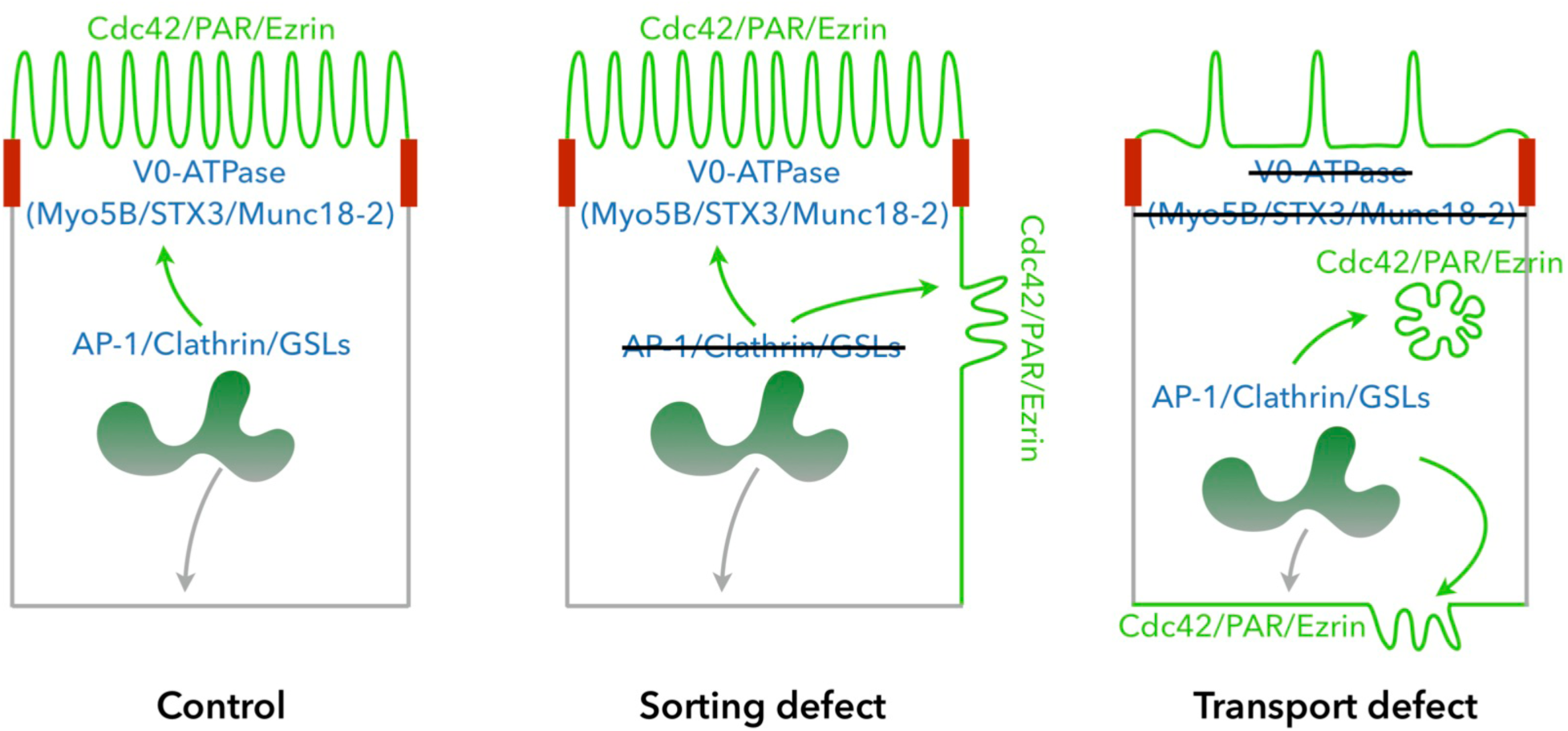
Comparison of polarity maintenance pathways. In control cells, AP-1, Clathrin and GSLs ensure the correct sorting of polarity and brush border proteins, directly or through adaptor proteins, to the apical PM. PM targeting likely involves RAB-11^+^ apical recycling endosomes, from which polarity/brush border/adaptors-containing vesicles fuse with the PM in a V0-ATPase/Myo5B/STX3/Munc18-2-dependent manner. Upon AP-1 and/or Clathrin and/or GSLs depletion, the early apical sorting step is defective, leading to a polarity inversion that induces the lateral accumulation of apical factors and the formation of lateraly positionned ectopic apical membranes. Upon V0-ATPase or Myo5B/STX3/Munc18-2 depletion, a late apical exocytosis step is defective, which affects only trafficking (i.e. SNAP-29 or STX3) and induces the misrouting of polarity/brush border components to the basolateral PM or in MVIs.

**Table S1.**
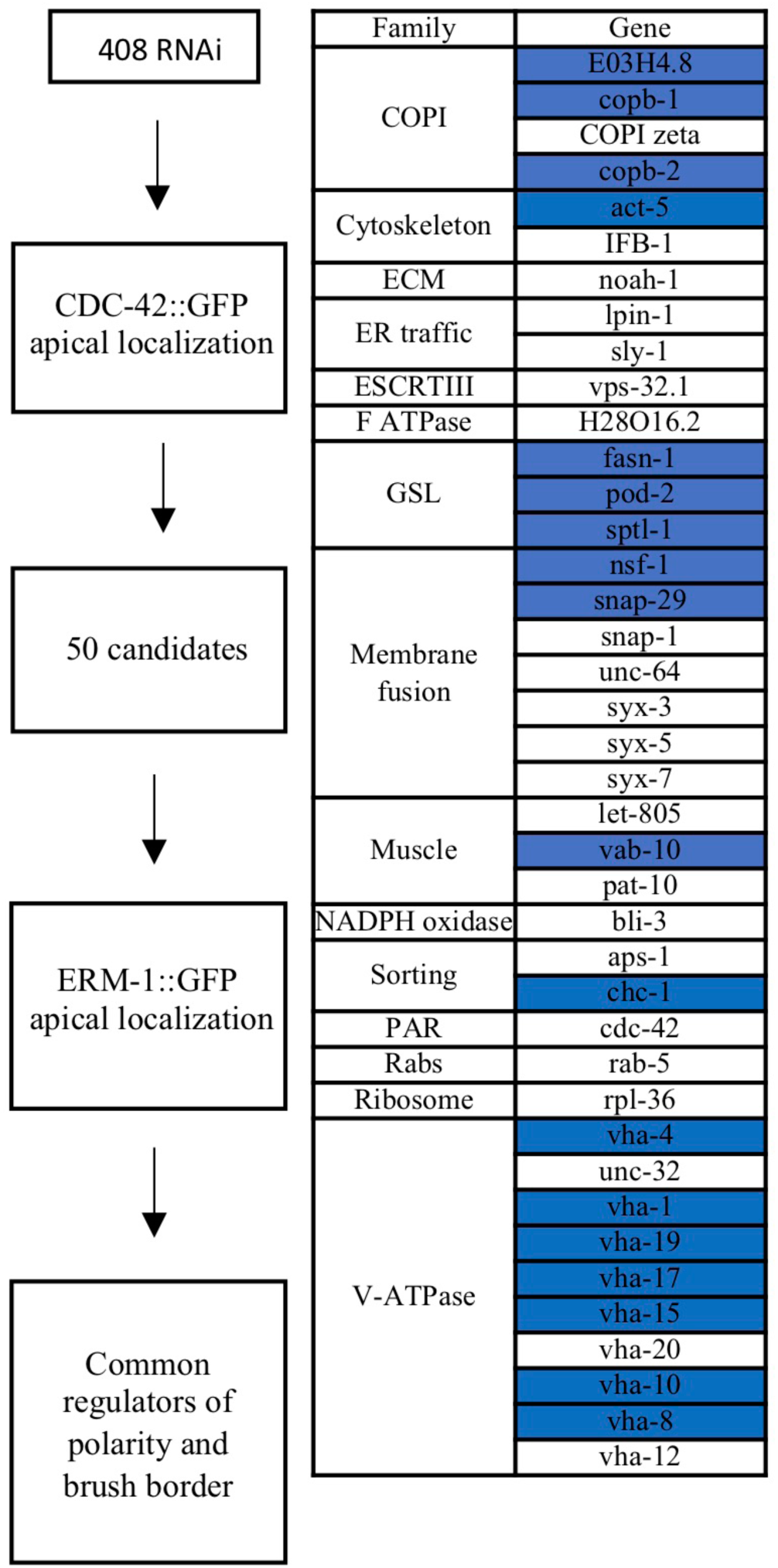
Polarity maintenance screen results. White and blue cases indicate the genes that affect CDC-42 only or CDC-42 and ERM-1 apical localization, respectively.

**Table S2.** Comparison of *vha-1* and *vha-8* silencing phenotypes.

|  | **L4440** | ***vha-1(RNAi) (V0)*** | ***vha-8(RNAi) (V1)*** |
| --- | --- | --- | --- |
| PAR-6, PKC-3, ERM-1<br>apical PM | Normal | Down | Down |
| Basolateral PAR-6, PKC-3,<br>ERM-1 | No | Yes | No |
| SNAP-29 cytoplasmic<br>accumulation | No | Yes | No |
| Mixed organelles | No | Yes | Yes |
| LMP-1, RAB-7, RAB-10<br>accumulation | No | Yes | Yes |
| Autophagy activation | No | +++ | + |
| Accumulation of<br>cytoplasmic lucent vesicles | No | Yes | No |
| RAB-11 signal | - | Down | Up |
| Microvillus atrophy | No | Yes | No |
| MVIs, vacuoles | No | Yes | No |

**Table S3.**
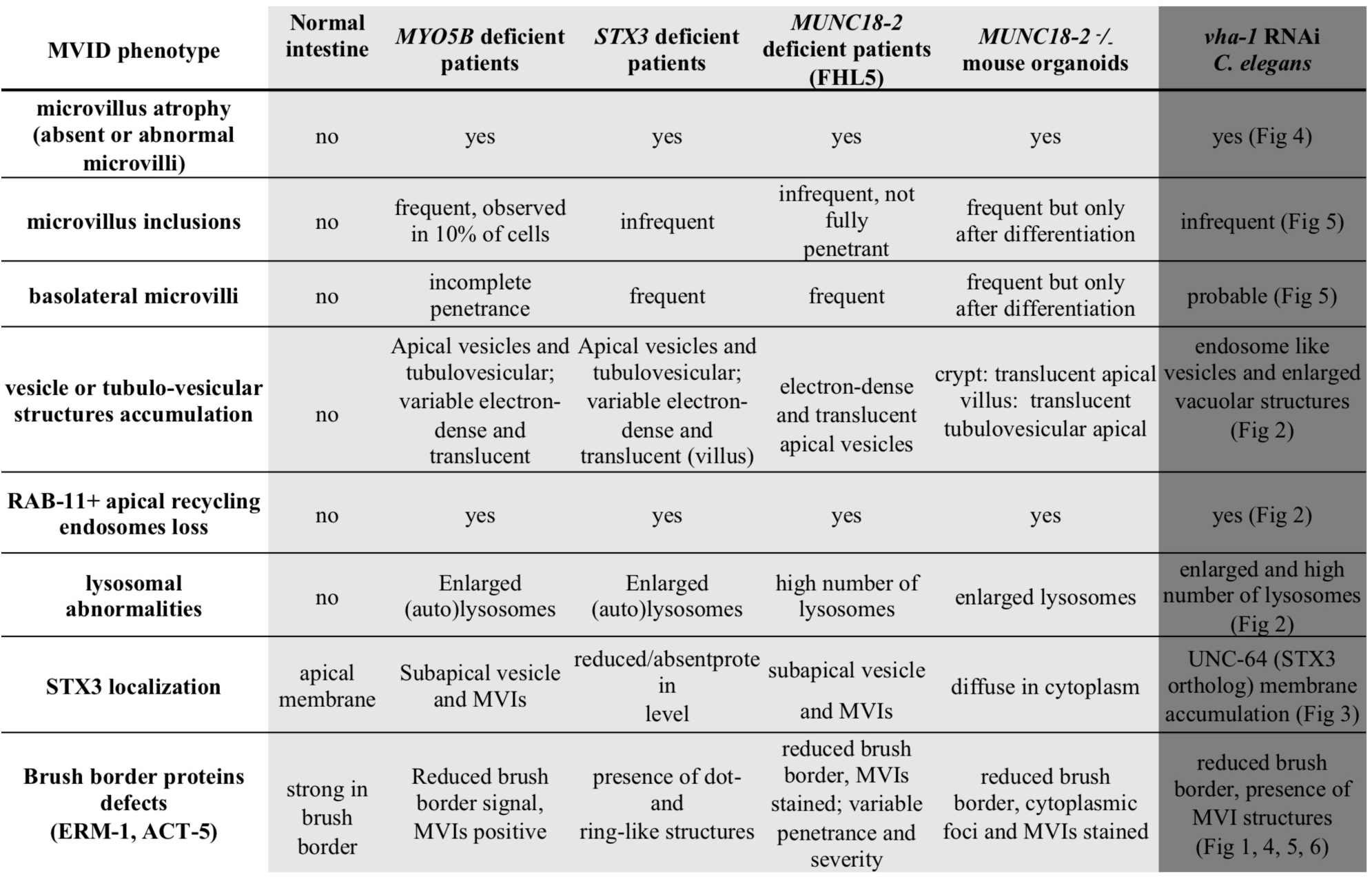
Comparison of MVID and V0-ATPase depletion phenotypes. Adapted from (49), with minor modifications.

**Table S4.**
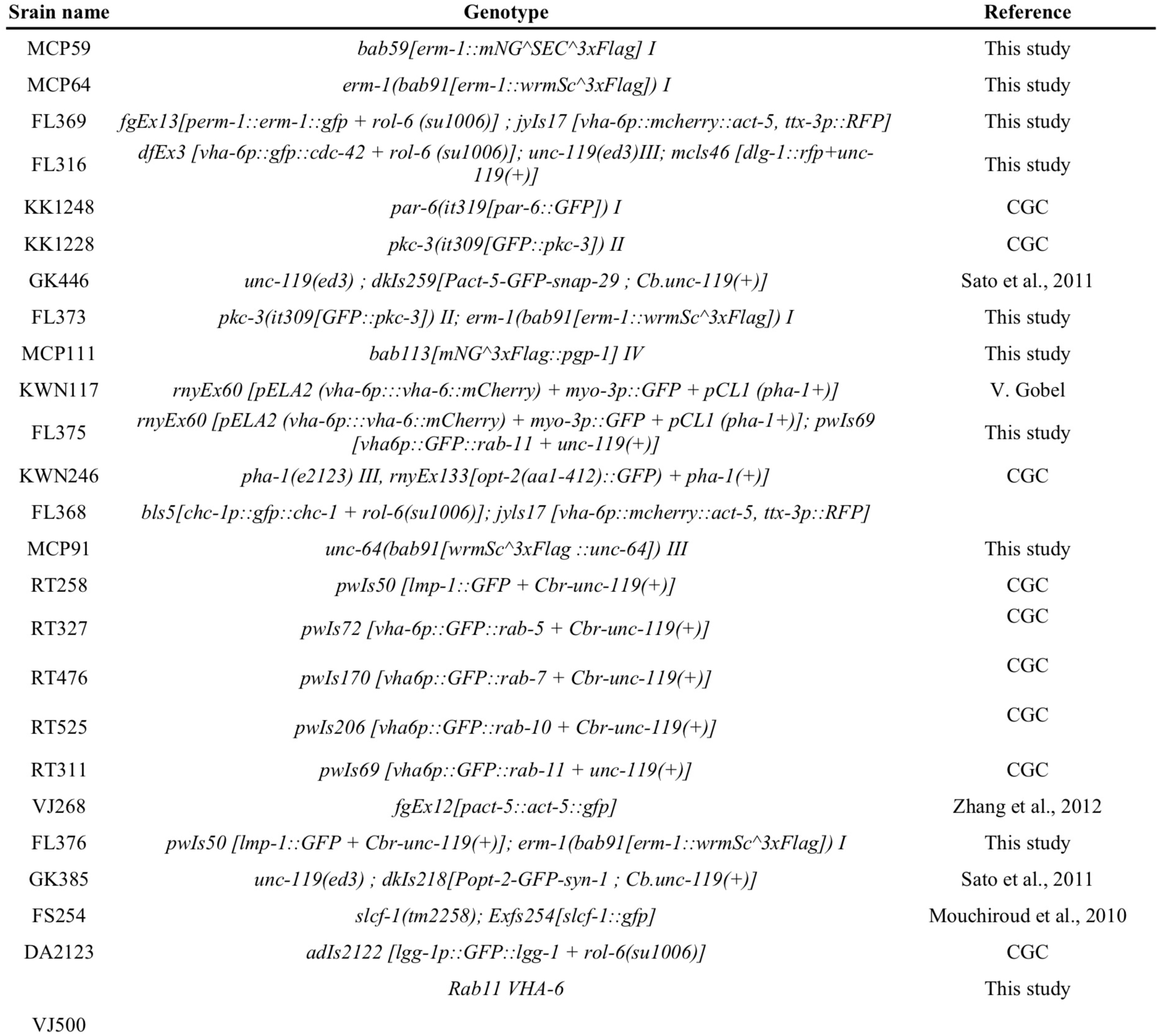
*C*. *elegans* strains used in this study.

**Movie S1. Cytoplasmic nucleation of MVIs.** Maximum projection of ERM-1::mNG expressing worms silenced for *vha-1* during 96h. Arrows indicate nucleating MVIs.

**Movie S2. Cytoplasmic nucleation of MVIs.** Maximum projection of ERM-1::mNG expressing worms silenced for *vha-1* during 96h. Arrowheads and arrows indicate irregular MVIs/vacuoles and cytoplasmic nucleating MVIs, respectively.

**Movie S3. MVI formation by internalization of the apical PM.** 3D projection of ERM-1::GFP expressing worms silenced for *vha-1* during 72h.

**Movie S4. MVI elimination.** Maximum projection of ERM-1::mNG expressing worms silenced for *vha-1* during 96h. The arrow indicates the elimination of the MVIs inside the presumptive autophagic ERM-1^+^ membrane. Arrowheads show direct MVI disappearance in the cytoplasm.

## References

Allman, E., Johnson, D. & Nehrke, K. 2009. Loss of the apical V-ATPase a-subunit VHA-6 prevents acidification of the intestinal lumen during a rhythmic behavior in C. elegans. Am J Physiol Cell Physiol, 297, C1071–81.

Baars, T. L., Petri, S., Peters, C. & Mayer, A. 2007. Role of the V-ATPase in regulation of the vacuolar fission-fusion equilibrium. Mol Biol Cell, 18, 3873–82.

Balklava, Z., Pant, S., Fares, H. & Grant, B. D. 2007. Genome-wide analysis identifies a general requirement for polarity proteins in endocytic traffic. Nat Cell Biol, 9, 1066–73.

Bonifacino, J. S. 2014. Adaptor proteins involved in polarized sorting. J Cell Biol, 204, 7–17.

Brenner, S. 1974. The genetics of Caenorhabditis elegans. Genetics, 77, 71–94.

Bryant, D. M., Datta, A., Rodriguez-Fraticelli, A. E., Peranen, J., Martin-Belmonte, F. & Mostov, K. E. 2010. A molecular network for de novo generation of the apical surface and lumen. Nat Cell Biol, 12, 1035–45.

Chang, J. T., Kumsta, C., Hellman, A. B., Adams, L. M. & Hansen, M. 2017. Spatiotemporal regulation of autophagy during Caenorhabditis elegans aging. Elife, 6.

Collaco, A. M., Geibel, P., Lee, B. S., Geibel, J. P. & Ameen, N. A. 2013. Functional vacuolar ATPase (V-ATPase) proton pumps traffic to the enterocyte brush border membrane and require CFTR. Am J Physiol Cell Physiol, 305, C981–96.

Colombie, N., Choesmel-Cadamuro, V., Series, J., Emery, G., Wang, X. & Ramel, D. 2017. Non-autonomous role of Cdc42 in cell-cell communication during collective migration. Dev Biol, 423, 12–18.

Crawley, S. W., Mooseker, M. S. & Tyska, M. J. 2014. Shaping the intestinal brush border. J Cell Biol, 207, 441–51.

Desclozeaux, M., Venturato, J., Wylie, F. G., Kay, J. G., Joseph, S. R., Le, H. T. & Stow, J. L. 2008. Active Rab11 and functional recycling endosome are required for E-cadherin trafficking and lumen formation during epithelial morphogenesis. Am J Physiol Cell Physiol, 295, C545–56.

Dhekne, H. S., Hsiao, N. H., Roelofs, P., Kumari, M., Slim, C. L., Rings, E. H. & Van Ijzendoorn, S. C. 2014. Myosin Vb and Rab11a regulate phosphorylation of ezrin in enterocytes. J Cell Sci, 127, 1007–17.

Dhekne, H. S., Pylypenko, O., Overeem, A. W., Ferreira, R. J., Van Der Velde, K. J., Rings, E., Posovszky, C., Swertz, M. A., Houdusse, A. & Van, I. S. C. D. 2018. MYO5B, STX3, and STXBP2 mutations reveal a common disease mechanism that unifies a subset of congenital diarrheal disorders: A mutation update. Hum Mutat, 39, 333–344.

Di Giovanni, J., Boudkkazi, S., Mochida, S., Bialowas, A., Samari, N., Leveque, C., Youssouf, F., Brechet, A., Iborra, C., Maulet, Y., Moutot, N., Debanne, D., Seagar, M. & El Far, O. 2010. V-ATPase membrane sector associates with synaptobrevin to modulate neurotransmitter release. Neuron, 67, 268–79.

Engevik, A. C., Kaji, I., Engevik, M. A., Meyer, A. R., Weis, V. G., Goldstein, A., Hess, M. W., Muller, T., Koepsell, H., Dudeja, P. K., Tyska, M., Huber, L. A., Shub, M. D., Ameen, N. & Goldenring, J. R. 2018. Loss of MYO5B Leads to Reductions in Na(+) Absorption With Maintenance of CFTR-Dependent Cl(-) Secretion in Enterocytes. Gastroenterology.

Feng, Q., Bonder, E. M., Engevik, A. C., Zhang, L., Tyska, M. J., Goldenring, J. R. & Gao, N. 2017. Disruption of Rab8a and Rab11a causes formation of basolateral microvilli in neonatal enteropathy. J Cell Sci, 130, 2491–2505.

Forgac, M. 2007. Vacuolar ATPases: rotary proton pumps in physiology and pathophysiology. Nat Rev Mol Cell Biol, 8, 917–29.

Gillard, G., Shafaq-Zadah, M., Nicolle, O., Damaj, R., Pecreaux, J. & Michaux, G. 2015. Control of E-cadherin apical localisation and morphogenesis by a SOAP-1/AP-1/clathrin pathway in C. elegans epidermal cells. Development, 142, 1684–94.

Gravotta, D., Carvajal-Gonzalez, J. M., Mattera, R., Deborde, S., Banfelder, J. R., Bonifacino, J. S. & Rodriguez-Boulan, E. 2012. The Clathrin Adaptor AP-1A Mediates Basolateral Polarity. Dev Cell, 22, 811–23.

Guo, B., Liang, Q., Li, L., Hu, Z., Wu, F., Zhang, P., Ma, Y., Zhao, B., Kovacs, A. L., Zhang, Z., Feng, D., Chen, S. & Zhang, H. 2014. O-GlcNAc-modification of SNAP-29 regulates autophagosome maturation. Nat Cell Biol, 16, 1215–26.

Halac, U., Lacaille, F., Joly, F., Hugot, J. P., Talbotec, C., Colomb, V., Ruemmele, F. M. & Goulet, O. 2011. Microvillous inclusion disease: how to improve the prognosis of a severe congenital enterocyte disorder. J Pediatr Gastroenterol Nutr, 52, 460–5.

Hase, K., Nakatsu, F., Ohmae, M., Sugihara, K., Shioda, N., Takahashi, D., Obata, Y., Furusawa, Y., Fujimura, Y., Yamashita, T., Fukuda, S., Okamoto, H., Asano, M., Yonemura, S. & Ohno, H. 2013. AP-1B-Mediated Protein Sorting Regulates Polarity and Proliferation of Intestinal Epithelial Cells in Mice. Gastroenterology, 145, 625–35.

Hegan, P. S., Giral, H., Levi, M. & Mooseker, M. S. 2012. Myosin VI is required for maintenance of brush border structure, composition, and membrane trafficking functions in the intestinal epithelial cell. Cytoskeleton (Hoboken), 69, 235–51.

Hein, Z., Schmidt, S., Zimmer, K. P. & Naim, H. Y. 2011. The dual role of annexin II in targeting of brush border proteins and in intestinal cell polarity. Differentiation, 81, 243–52.

Heintzelman, M. B. & Mooseker, M. S. 1990a. Assembly of the brush border cytoskeleton: changes in the distribution of microvillar core proteins during enterocyte differentiation in adult chicken intestine. Cell Motil Cytoskeleton, 15, 12–22.

Heintzelman, M. B. & Mooseker, M. S. 1990b. Structural and compositional analysis of early stages in microvillus assembly in the enterocyte of the chick embryo. Differentiation, 43, 175–82.

Hiesinger, P. R., Fayyazuddin, A., Mehta, S. Q., Rosenmund, T., Schulze, K. L., Zhai, R. G., Verstreken, P., Cao, Y., Zhou, Y., Kunz, J. & Bellen, H. J. 2005. The v-ATPase V0 subunit a1 is required for a late step in synaptic vesicle exocytosis in Drosophila. Cell, 121, 607–20.

Iancu, T. C., Mahajnah, M., Manov, I. & Shaoul, R. 2007. Microvillous inclusion disease: ultrastructural variability. Ultrastruct Pathol, 31, 173–88.

Ji, Y. J., Choi, K. Y., Song, H. O., Park, B. J., Yu, J. R., Kagawa, H., Song, W. K. & Ahnn, J. 2006. VHA-8, the E subunit of V-ATPase, is essential for pH homeostasis and larval development in C. elegans. FEBS Lett, 580, 3161–6.

Kamath, R. S. & Ahringer, J. 2003. Genome-wide RNAi screening in Caenorhabditis elegans. Methods, 30, 313–21.

Kang, J., Bai, Z., Zegarek, M. H., Grant, B. D. & Lee, J. 2011. Essential roles of snap-29 in C. elegans. Dev Biol, 355, 77–88.

Kang, J., Shin, D., Yu, J. R. & Lee, J. 2009. Lats kinase is involved in the intestinal apical membrane integrity in the nematode Caenorhabditis elegans. Development, 136, 2705–15.

Kiela, P. R. & Ghishan, F. K. 2016. Physiology of Intestinal Absorption and Secretion. Best Pract Res Clin Gastroenterol, 30, 145–59.

Knight, A. J., Johnson, N. M. & Behm, C. A. 2012. VHA-19 is essential in Caenorhabditis elegans oocytes for embryogenesis and is involved in trafficking in oocytes. PLoS One, 7, e40317.

Knowles, B. C., Roland, J. T., Krishnan, M., Tyska, M. J., Lapierre, L. A., Dickman, P. S., Goldenring, J. R. & Shub, M. D. 2014. Myosin Vb uncoupling from RAB8A and RAB11A elicits microvillus inclusion disease. J Clin Invest, 124, 2947–62.

Kolotuev, I., Bumbarger, D. J., Labouesse, M. & Schwab, Y. 2012. Targeted ultramicrotomy: a valuable tool for correlated light and electron microscopy of small model organisms. Methods Cell Biol, 111, 203–22.

Kravtsov, D. V., Ahsan, M. K., Kumari, V., Van Ijzendoorn, S. C., Reyes-Mugica, M., Kumar, A., Gujral, T., Dudeja, P. K. & Ameen, N. A. 2016. Identification of intestinal ion transport defects in microvillus inclusion disease. Am J Physiol Gastrointest Liver Physiol, 311, G142–55.

Lee, S. K., Li, W., Ryu, S. E., Rhim, T. & Ahnn, J. 2010. Vacuolar (H+)-ATPases in Caenorhabditis elegans: what can we learn about giant H+ pumps from tiny worms? Biochim Biophys Acta, 1797, 1687–95.

Liegeois, S., Benedetto, A., Garnier, J. M., Schwab, Y. & Labouesse, M. 2006. The V0-ATPase mediates apical secretion of exosomes containing Hedgehog-related proteins in Caenorhabditis elegans. J Cell Biol, 173, 949–61.

Marion, S., Hoffmann, E., Holzer, D., Le Clainche, C., Martin, M., Sachse, M., Ganeva, I., Mangeat, P. & Griffiths, G. 2011. Ezrin promotes actin assembly at the phagosome membrane and regulates phagolysosomal fusion. Traffic, 12, 421– 37.

Martin-Belmonte, F., Gassama, A., Datta, A., Yu, W., Rescher, U., Gerke, V. & Mostov, K. 2007. PTEN-mediated apical segregation of phosphoinositides controls epithelial morphogenesis through Cdc42. Cell, 128, 383–97.

Maxson, M. E. & Grinstein, S. 2014. The vacuolar-type H(+)-ATPase at a glance - more than a proton pump. J Cell Sci, 127, 4987–93.

Mcghee, J. D. 2007. The C. elegans intestine. WormBook, 1–36.

Merkulova, M., Paunescu, T. G., Azroyan, A., Marshansky, V., Breton, S. & Brown, D. 2015. Mapping the H(+) (V)-ATPase interactome: identification of proteins involved in trafficking, folding, assembly and phosphorylation. Sci Rep, 5, 14827.

Michaux, G., Massey-Harroche, D., Nicolle, O., Rabant, M., Brousse, N., Goulet, O., Le Bivic, A. & Ruemmele, F. M. 2016. The localisation of the apical Par/Cdc42 polarity module is specifically affected in microvillus inclusion disease. Biol Cell, 108, 19–28.

Morel, N., Dedieu, J. C. & Philippe, J. M. 2003. Specific sorting of the a1 isoform of the V-H+ATPase a subunit to nerve terminals where it associates with both synaptic vesicles and the presynaptic plasma membrane. J Cell Sci, 116, 4751–62.

Mosa, M. H., Nicolle, O., Maschalidi, S., Sepulveda, F. E., Bidaud-Meynard, A., Menche, C., Michels, B. E., Michaux, G., De Saint Basile, G. & Farin, H. F. 2018. Dynamic Formation of Microvillus Inclusions During Enterocyte Differentiation in Munc18-2-Deficient Intestinal Organoids. Cell Mol Gastroenterol Hepatol, 6, 477–493 e1.

Muller, T., Hess, M. W., Schiefermeier, N., Pfaller, K., Ebner, H. L., Heinz-Erian, P., Ponstingl, H., Partsch, J., Rollinghoff, B., Kohler, H., Berger, T., Lenhartz, H., Schlenck, B., Houwen, R. J., Taylor, C. J., Zoller, H., Lechner, S., Goulet, O., Utermann, G., Ruemmele, F. M., Huber, L. A. & Janecke, A. R. 2008. MYO5B mutations cause microvillus inclusion disease and disrupt epithelial cell polarity. Nat Genet, 40, 1163–5.

Nicolle, O., Burel, A., Griffiths, G., Michaux, G. & Kolotuev, I. 2015. Adaptation of Cryo-Sectioning for IEM Labeling of Asymmetric Samples: A Study Using Caenorhabditis elegans. Traffic, 16, 893–905.

Reinshagen, K., Naim, H. Y. & Zimmer, K. P. 2002. Autophagocytosis of the apical membrane in microvillus inclusion disease. Gut, 51, 514–21.

Rodriguez-Boulan, E. & Macara, I. G. 2014. Organization and execution of the epithelial polarity programme. Nat Rev Mol Cell Biol, 15, 225–42.

Ruemmele, F. M., Schmitz, J. & Goulet, O. 2006. Microvillous inclusion disease (microvillous atrophy). Orphanet J Rare Dis, 1, 22.

Saegusa, K., Sato, M., Sato, K., Nakajima-Shimada, J., Harada, A. & Sato, K. 2014. Caenorhabditis elegans chaperonin CCT/TRiC is required for actin and tubulin biogenesis and microvillus formation in intestinal epithelial cells. Mol Biol Cell, 25, 3095–104.

Sato, M., Saegusa, K., Sato, K., Hara, T., Harada, A. & Sato, K. 2011. Caenorhabditis elegans SNAP-29 is required for organellar integrity of the endomembrane system and general exocytosis in intestinal epithelial cells. Mol Biol Cell, 22, 2579–87.

Schneeberger, K., Roth, S., Nieuwenhuis, E. E. S. & Middendorp, S. 2018. Intestinal epithelial cell polarity defects in disease: lessons from microvillus inclusion disease. Dis Model Mech, 11.

Schneeberger, K., Vogel, G. F., Teunissen, H., Van Ommen, D. D., Begthel, H., El Bouazzaoui, L., Van Vugt, A. H., Beekman, J. M., Klumperman, J., Muller, T., Janecke, A., Gerner, P., Huber, L. A., Hess, M. W., Clevers, H., Van Es, J. H., Nieuwenhuis, E. E. & Middendorp, S. 2015. An inducible mouse model for microvillus inclusion disease reveals a role for myosin Vb in apical and basolateral trafficking. Proc Natl Acad Sci U S A, 112, 12408–13.

Shafaq-Zadah, M., Brocard, L., Solari, F. & Michaux, G. 2012. AP-1 is required for the maintenance of apico-basal polarity in the C. elegans intestine. Development, 139, 2061–70.

Sidhaye, J., Pinto, C. S., Dharap, S., Jacob, T., Bhargava, S. & Sonawane, M. 2016. The zebrafish goosepimples/myosin Vb mutant exhibits cellular attributes of human microvillus inclusion disease. Mech Dev, 142, 62–74.

Sobajima, T., Yoshimura, S., Iwano, T., Kunii, M., Watanabe, M., Atik, N., Mushiake, S., Morii, E., Koyama, Y., Miyoshi, E. & Harada, A. 2014. Rab11a is required for apical protein localisation in the intestine. Biol Open, 4, 86–94.

Sobota, J. A., Back, N., Eipper, B. A. & Mains, R. E. 2009. Inhibitors of the V0 subunit of the vacuolar H+-ATPase prevent segregation of lysosomal- and secretory-pathway proteins. J Cell Sci, 122, 3542–53.

Steegmaier, M., Yang, B., Yoo, J. S., Huang, B., Shen, M., Yu, S., Luo, Y. & Scheller, R. H. 1998. Three novel proteins of the syntaxin/SNAP-25 family. J Biol Chem, 273, 34171–9.

Sudhof, T. C. & Rothman, J. E. 2009. Membrane fusion: grappling with SNARE and SM proteins. Science, 323, 474–7.

Talmon, G., Holzapfel, M., Dimaio, D. J. & Muirhead, D. 2012. Rab11 is a useful tool for the diagnosis of microvillous inclusion disease. Int J Surg Pathol, 20, 252–6.

Vacca, B., Bazellieres, E., Nouar, R., Harada, A., Massey-Harroche, D. & Le Bivic, A. 2014. Drebrin E depletion in human intestinal epithelial cells mimics Rab8a loss of function. Hum Mol Genet, 23, 2834–46.

Vogel, G. F., Hess, M. W., Pfaller, K., Huber, L. A., Janecke, A. R. & Muller, T. 2016. Towards understanding microvillus inclusion disease. Mol Cell Pediatr, 3, 3.

Vogel, G. F., Janecke, A. R., Krainer, I. M., Gutleben, K., Witting, B., Mitton, S. G., Mansour, S., Ballauff, A., Roland, J. T., Engevik, A. C., Cutz, E., Muller, T., Goldenring, J. R., Huber, L. A. & Hess, M. W. 2017a. Abnormal Rab11-Rab8-vesicles cluster in enterocytes of patients with microvillus inclusion disease. Traffic, 18, 453–464.

Vogel, G. F., Klee, K. M., Janecke, A. R., Muller, T., Hess, M. W. & Huber, L. A. 2015. Cargo-selective apical exocytosis in epithelial cells is conducted by Myo5B, Slp4a, Vamp7, and Syntaxin 3. J Cell Biol, 211, 587–604.

Vogel, G. F., Van Rijn, J. M., Krainer, I. M., Janecke, A. R., Posovszky, C., Cohen, M., Searle, C., Jantchou, P., Escher, J. C., Patey, N., Cutz, E., Muller, T., Middendorp, S., Hess, M. W. & Huber, L. A. 2017b. Disrupted apical exocytosis of cargo vesicles causes enteropathy in FHL5 patients with Munc18-2 mutations. JCI Insight, 2.

Weis, V. G., Knowles, B. C., Choi, E., Goldstein, A. E., Williams, J. A., Manning, E. H., Roland, J. T., Lapierre, L. A. & Goldenring, J. R. 2016. Loss of MYO5B in mice recapitulates Microvillus Inclusion Disease and reveals an apical trafficking pathway distinct to neonatal duodenum. Cell Mol Gastroenterol Hepatol, 2, 131–157.

Wiegerinck, C. L., Janecke, A. R., Schneeberger, K., Vogel, G. F., Van Haaften-Visser, D. Y., Escher, J. C., Adam, R., Thoni, C. E., Pfaller, K., Jordan, A. J., Weis, C. A., Nijman, I. J., Monroe, G. R., Van Hasselt, P. M., Cutz, E., Klumperman, J., Clevers, H., Nieuwenhuis, E. E., Houwen, R. H., Van Haaften, G., Hess, M. W., Huber, L. A., Stapelbroek, J. M., Muller, T. & Middendorp, S. 2014. Loss of syntaxin 3 causes variant microvillus inclusion disease. Gastroenterology, 147, 65–68 e10.

Winter, J. F., Hopfner, S., Korn, K., Farnung, B. O., Bradshaw, C. R., Marsico, G., Volkmer, M., Habermann, B. & Zerial, M. 2012. Caenorhabditis elegans screen reveals role of PAR-5 in RAB-11-recycling endosome positioning and apicobasal cell polarity. Nat Cell Biol, 14, 666–76.

Zhang, H., Abraham, N., Khan, L. A., Hall, D. H., Fleming, J. T. & Gobel, V. 2011. Apicobasal domain identities of expanding tubular membranes depend on glycosphingolipid biosynthesis. Nat Cell Biol, 13, 1189–201.

Zhang, H., Kim, A., Abraham, N., Khan, L. A., Hall, D. H., Fleming, J. T. & Gobel, V. 2012. Clathrin and AP-1 regulate apical polarity and lumen formation during C. elegans tubulogenesis. Development, 139, 2071–83.

Zhu, H., Sewell, A. K. & Han, M. 2015. Intestinal apical polarity mediates regulation of TORC1 by glucosylceramide in C. elegans. Genes Dev, 29, 1218–23.

